# Integrated analysis of glycan and RNA in single cells

**DOI:** 10.1101/2020.06.15.153536

**Authors:** Fumi Minoshima, Haruka Ozaki, Haruki Odaka, Hiroaki Tateno

## Abstract

Single-cell sequencing has emerged as an indispensable technology to dissect cellular heterogeneity but never been applied to the simultaneous analysis of glycan and RNA. Using oligonucleotide-labeled lectins, we first established lectin-based glycan profiling of single cells by sequencing (scGlycan-seq). We then combined the scGlycan-seq with single-cell transcriptome profiling (scRNA-seq) for joint analysis of glycan and RNA in single cells (scGR-seq). Using scGR-seq, we analyzed the two modalities in human induced pluripotent stem cells (hiPSCs) before and after differentiation into neural progenitor cells at the single cell resolution. The combination of RNA and glycan separated the two cell types clearer than either one of them. Furthermore, integrative analysis of glycan and RNA modalities in single cells found known and novel lectins that were specific to hiPSCs and coordinated with neural differentiation. Taken together, we demonstrate that scGR-seq can reveal the cellular heterogeneity and biological roles of glycans across multicellular systems.

## Introduction

Glycans present at the surface of all living cells play crucial roles in diverse biological processes. Glycan structures have been known to vary depending on cell types and states. Therefore, glycans are often referred to as “cell signature” that reflect cellular characteristics. Indeed, most of the stem cell markers^1–5^ and serum tumor markers are glycans^6–8^. Glycans are the secondary products of genes synthesized by the orchestration of many proteins, such as glycosyltransferases and glycosidases. There has been no method to predict the precise structure of the complex glycans only from the gene expression profiles. Therefore, it is vital to develop methods to analyze cell surface glycans directly. In this sense, different strategies have been undertaken to analyze glycome, including mass spectrometry (MS), HPLC, nuclear magnetic resonance (NMR), and capillary electrophoresis (CE)^9–11^. Recently, a lectin-based glycan profiling technology called lectin microarray has played a pivotal role in surveying and mapping the informational context of complex glycans of various biological samples, indicating the applicability of lectin-based glycan profiling^12–14^. However, there are limitations to the current glycan analytical methods. For instance, (1) glycans are unable to be analyzed at a single-cell level, (2) the glycan profile of each cell type in the mixed cell populations cannot be obtained without prior cell separation, and (3) the relationship between glycome and transcriptome in single cells cannot be analyzed. Simultaneous analysis of the two modalities in single cells could lead to the understanding of cellular heterogeneity and glycan functions, and the development of novel glycan markers of rare cells.

High-throughput single-cell sequencing has been transformative to understand the complex cell populations^15^. Recently, simultaneous profiling of multiple types of molecules within a single cell has been developed for building a much more comprehensive molecular view of the cell^15–17^. However, there has been no technology to jointly analyze glycome and transcriptome in single cells, since glycans cannot be amplified by PCR, unlike DNA and RNA. Here, we first established highly-multiplexed lectin-based glycan profiling of single cells by sequencing (scGlycan-seq). We then combined the scGlycan-seq with single-cell transcriptome profiling (scRNA-seq) for joint analysis of glycan and RNA in single cells (scGR-seq). Using scGR-seq, we analyzed the relationship between the two distinct layers in human induced pluripotent stem cells (hiPSCs) before and after differentiation into neural progenitor cells (NPCs) in single cells and revealed the cellular heterogeneity across mRNA and glycans even within cells of the same cell types.

## Results

### Principle of Glycan-seq

We hypothesized that a lectin conjugated with a DNA oligonucleotide (DNA-barcoded lectin) could be measured by sequencing as a digital readout of glycan abundance^18^. To address this possibility, we conjugated lectins to DNA oligonucleotides designed to contain a barcode sequence for the identification of lectin and allowed specific identification by PCR (Fig. 1a). Lectins were conjugated via their amino groups with photocleavable dibenzocyclooctyne-N-hydroxysuccinimidyl ester (DBCO-NHS), which allowed efficient conjugation with 5’-azide-modified oligonucleotides (Supplementary Fig. 1a). The lectin-to-oligonucleotide ratio was confirmed by measuring the concentration of DNA and lectin (Supplementary Table 1). The oligonucleotides were released from the lectin by 15 min of ultraviolet (UV) exposure. Since the liberated amount of DNA increased as the UV exposure time by qRT-PCR (Supplementary Fig. 2), we determined the UV exposure time to balance the liberated amount of DNA and cell damage by UV exposure^19^. DNA-barcoded lectins were then purified by affinity chromatography to remove excess DNA oligonucleotides. We prepared a panel of 39 DNA-barcoded lectins that exhibited distinct glycan-binding specificities, and DNA-barcoded mouse and goat IgG as negative controls (Supplementary Table 1). Total 41 DNA-barcoded proteins were incubated with 1×10^5^ cells and unbound lectins were removed by washing (Fig. 1b). Then, single-cell or 1×10^4^ cells were separated into PCR tubes and exposed to UV. After centrifugation, supernatants containing released DNA barcodes were recovered, amplified by PCR, and analyzed by next-generation sequencing (NGS) to count the DNA barcodes. We designated this method as Glycan-seq.

**Fig. 1:**
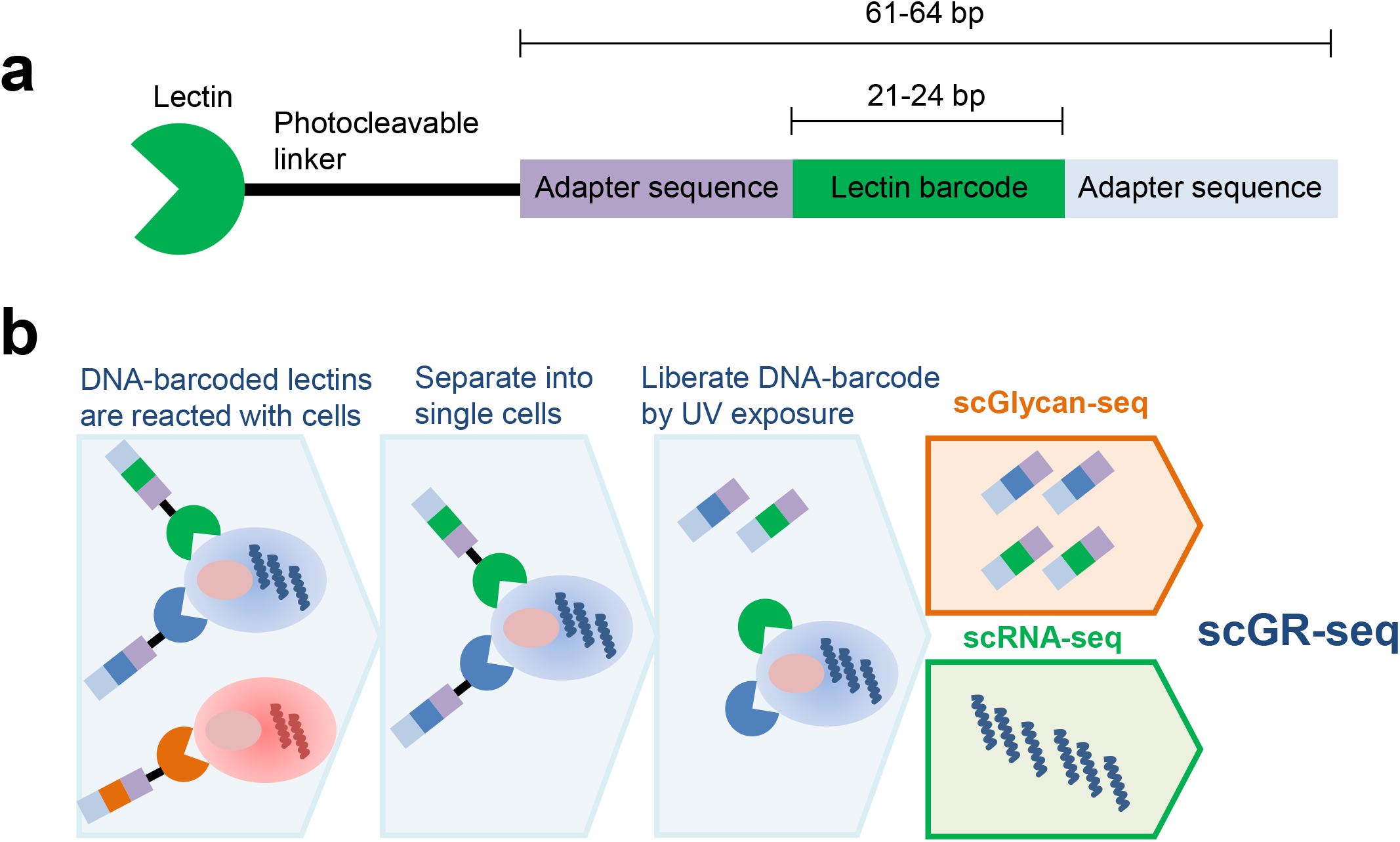
Principle of scGlycan-seq and scGR-seq. **(a)** Illustration of the DNA-barcoded lectin. **(b)** Schematic representation of scGlycan-seq, scRNA-seq, and scGR-seq. Cells were incubated with DNA-barcoded lectins and separated into single cells. After DNA-barcode was released by UV exposure, the released DNA-barcode in the cell culture supernatants was measured by next-generation sequencing. RNA transcripts were purified from the cell pellet and analyzed by scRNA-seq.

To evaluate the PCR amplification bias of DNA barcodes, we prepared a mix of equal amount of 41 DNA barcodes conjugated to 41 probes and performed Glycan-seq following 20 cycles of PCR. The barcode counts for each DNA barcode were divided by those of all DNA barcodes and expressed as percentage (%). The average percentage was 2.43% and the variation of the detected DNA barcodes was ranged between 0.26 and 1.58-fold of the average percentage (Supplementary Fig. 3), suggesting that the PCR bias is less than 2 PCR cycles. Therefore, we considered that PCR bias is within the allowable range.

### Glycan-seq of bulk cell populations

We assessed the ability of Glycan-seq to discriminate distinct cell populations based on cell surface glycan expression in bulk samples. Obtained data were compared with flow cytometry using fluorescence-labeled lectins as the gold standard.

We first applied Glycan-seq to hiPSCs and human dermal fibroblasts (hFibs) with triplicate. We focused on rBC2LCN, known to specifically bind to hiPSCs but not to hFibs^20–22^. A higher percentage of DNA-barcoded rBC2LCN was detected in hiPSCs (42.4%) than hFibs (0.4%), which was consistent with flow cytometry results (Figs. 2a and 2b, Supplementary Tables 2 and 3). Mouse and sheep IgG were used as negative controls, providing negligible DNA barcode levels (<0.01%) (Supplementary Tables 2 and 3). Glycan-seq data of other lectins such as rAAL, rAOL, TJAII, and rPALa also agreed well with flow cytometry data (Supplementary Fig. 4).

**Fig. 2:**
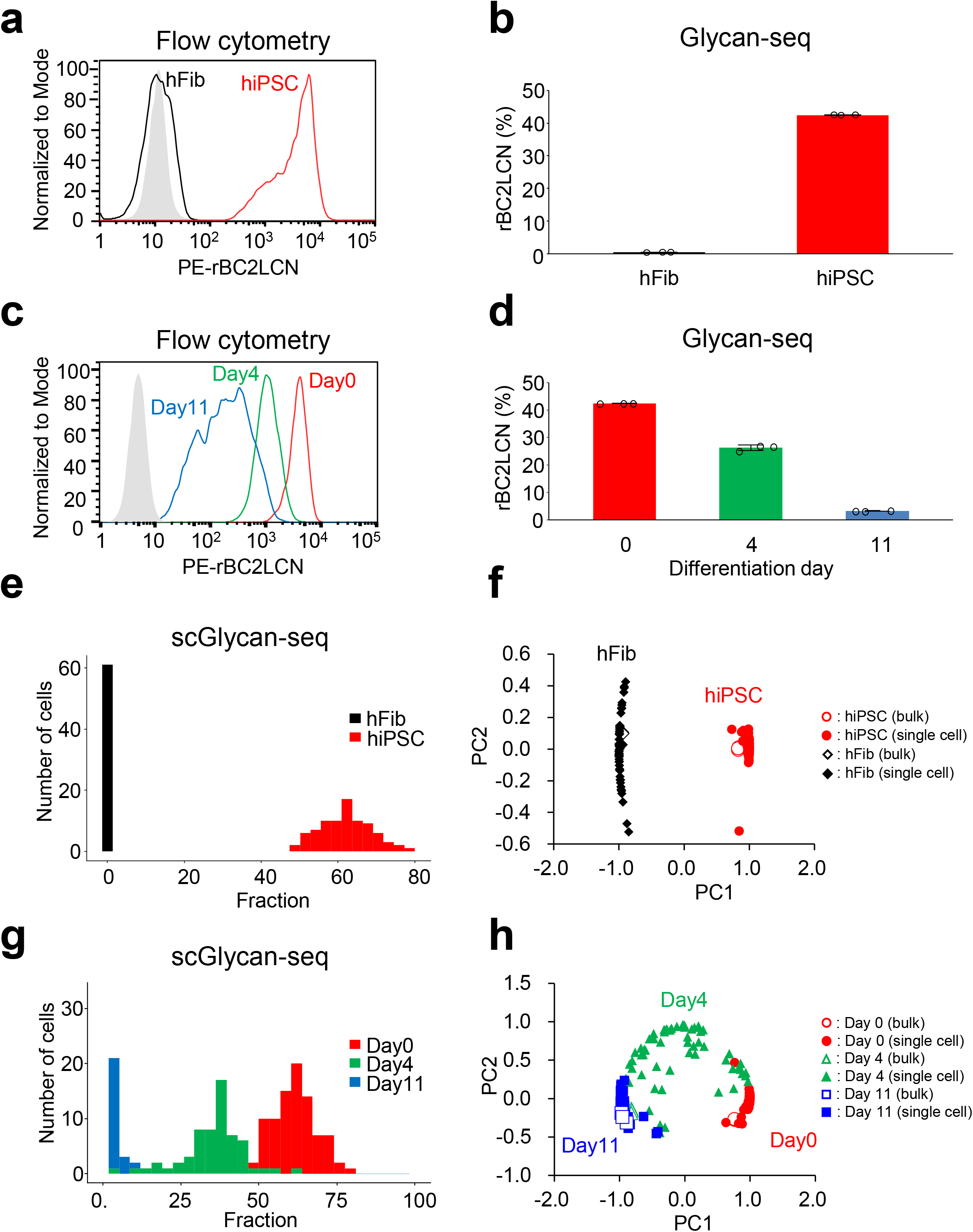
Comparison of Glycan-seq and flow cytometry. **(a)** Binding of R-phycoerythrin (PE)-labeled rBC2LCN to hiPSCs (*red line*) and hFibs (*black line*) was analyzed by flow cytometry. *Grey*: Binding of PE-labeled BSA to hiPSCs (negative control). **(b)** Binding of DNA-barcoded rBC2LCN to hiPSCs and hFibs was analyzed by Glycan-seq. The number of DNA-barcode derived from rBC2LCN was divided by that of the DNA-barcode of all lectins, multiplied by 100, and expressed as percentage (%). Data are shown as average ± SD of triplicates of the same sample. Each data was also plotted on a graph. **(c)** Binding of PE-labeled rBC2LCN to hiPSCs after differentiation to NPCs for 0 (*red line*), 4 (*green line*), and 11 days (*blue line*) was analyzed by flow cytometry. **(d)** Binding of DNA-barcoded rBC2LCN to hiPSCs after differentiation to NPCs for 0 (*red bar*), 4 (*green bar*), and 11 days (*blue bar*) was analyzed by Glycan-seq. Data are shown as average ± SD of triplicates of the same sample. Each data was also plotted on a graph. **(e)** Binding of DNA-barcoded rBC2LCN to hiPSCs and hFibs analyzed by scGlycan-seq. The histogram of rBC2LCN signal levels measured for hiPSCs (n = 83, *red*) and hFibs (n = 61, *black*) is shown. There was a statistically significant in the distribution of rBC2LCN between the two cell types (p < 0.001, Brunner-Munzel test with bonferroni correction). **(f)** PCA of bulk (100 cells, n = 3 for each cell type) and single cells (n = 96 for each cell type) of hiPSCs (*filled circle*: single cell, *open circle*: bulk) and hFibs (*filled triangle*: single cell, *open triangle*: bulk). PC1: principal component 1. PC2: principal component 2. **(g)** Binding of DNA-barcoded rBC2LCN to hiPSCs after differentiation to NPCs analyzed by scGlycan-seq. The histogram of rBC2LCN signal levels measured for hiPSCs after differentiation to NPCs for 0 (n = 84, *red*), 4 (n = 61, *blue*), 11 days (n = 57, *green*) is shown. There was a statistically significant in the distribution of rBC2LCN between the two cell types (Day0 vs Day4; p < 0.001, Day4 vs Day11; p < 0.001, Day0 vs Day11; p < 0.001, Brunner-Munzel test with bonferroni correction). **(h)** PCA of bulk (100 cells, n = 3 for each cell type) and single cells (n = 96 for each cell type) of hiPSCs after differentiation to NPCs for 0 (*open circle*: bulk, *filled circle*: single cell), 4 (*open triangle*: bulk, *filled triangle*: single cell), and 11 days (*open square*: bulk, *filled square*: single cell). DNA-barcode derived from each lectin was divided by DNA-barcode of all lectins and multiplied by 100. Glycan-seq data are available in Supplementary Tables 2, 3, 4, and 5. PC1: principal component 1. PC2: principal component 2.

We next applied Glycan-seq to Chinese hamster ovary (CHO) cells and glycosylation-defective mutants (Lec1 and Lec8) (Supplementary Fig. 5)^23^. Wild-type (WT) cells typically express complex-type N-glycans^24^, while Lec8 and Lec1 express agalactosylated and mannosylated N-glycans, respectively. In both flow cytometry and Glycan-seq, rLSLN (galactose binder) showed higher binding to WT than Lec8 and Lec1. Similarly, rSRL (GlcNAc binder) and rCalsepa (mannose binder) showed strong binding to Lec8 and Lec1, respectively. These results demonstrated that Glycan-seq could distinguish different bulk cell populations depending on cell surface glycan expression.

We further addressed whether relative quantitative differences in expression levels observed by flow cytometry could be measured by Glycan-seq. We applied Glycan-seq to hiPSCs before (Day 0) and after (Day 4 and 11) differentiation to NPCs, which were fluorescently-stained with NPC marker (NESTIN and PAX6) antibodies in fluorescence staining and flow cytometry (Supplementary Figs. 6 and 7). Using qRT-PCR, NPC marker genes (*SOX1*, *NESTIN*, *PAX6*, *FOXG1*), but not an hPSC marker gene (*OCT4*)^25,26^, were detected in NPCs (Supplementary Fig. 8), indicating that hiPSCs could be successfully differentiated into NPCs. In flow cytometry, rBC2LCN binding gradually decreased during differentiation into NPCs (Fig. 2c). Similar trends were observed for rBC2LCN signal in Glycan-seq (Fig. 2d, Supplementary Tables 2 and 3). Collectively, these results indicate that bulk Glycan-seq can capture distinct and quantitative differences in glycan profiles in various cell populations as observed by flow cytometry.

### Single-cell Glycan-seq

We then tested Glycan-seq to see its applicability in single cells, which we termed single-cell Glycan-seq (scGlycan-seq) (Fig. 1b). We first applied scGlycan-seq to hiPSCs and hFibs. To assess the effect of the total barcode counts in each cell on the obtained glycan profiles, we performed principal component analysis (PCA) on the scGlycan-seq data. Single cells of hFibs with low total barcode counts showed low values in PC1 (Supplementary Fig. 9a). Since the cells with low and high total barcode counts were separated in the histogram (Supplementary Fig. 9b), we determined the cut-off value of the total counts by Otsu’s method, and hFibs with higher than 19,465 total counts were selected and applied for further analyses (Supplementary Fig. 9c). The same quality control was adopted to hiPSCs, and hiPSCs with higher than 6,126 total counts were used for the following analysis.

The signal level of rBC2LCN in hiPSCs and hFibs obtained by scGlycan-seq was shown in Fig. 2e. scGlycan-seq recapitulated heterogeneity of rBC2LCN observed by flow cytometry and showed statistically significant differences between hiPSCs and hFibs (p < 0.001, Brunner-Munzel test). We then performed PCA on scGlycan-seq data together with bulk Glycan-seq data of hiPSCs and hFibs (Fig. 2f). The PC1 clearly separated the two cell types and, for each of hiPSCs and hiFibs, the PC2 showed higher variability of single cells compared to bulk samples, revealing cell-to-cell heterogeneity in glycan profiles (Fig.2f, Supplementary Tables 4 and 5).

We further applied scGlycan-seq to the hiPSCs after differentiation into NPCs (days 0, 4, and 11). The signal level of each lectin in hiPSCs and 11-day NPCs was shown in Supplementary Fig. 10. Relative quantitative differences in the rBC2LCN signal for hiPSCs before (Day 0) and after differentiation to NPCs (Day 4 and 11) observed by flow cytometry could also be captured by scGlycan-seq (Fig. 2c, g). Statistically significant differences between hiPSCs (Day 0) and hiPSC-derived NPCs (Day 4 and11) (p < 0.001, Brunner-Munzel test) were observed. PCA clearly separated single cells of Day 0, 4, and 11, and cells were ordered as differentiation progression (Fig. 2h). Single-cell heterogeneity of hiPSCs increased after 4-day differentiation, but converged after 11-day differentiation, possibly because 4-day NPCs might contain various degrees of differentiated NPCs (Fig. 2h, Supplementary Tables 4 and 5). These results demonstrated that scGlycan-seq enabled glycan profiling in single cells and revealed cellular heterogeneity in glycan profiles.

### scGR-seq of hiPSCs and NPCs

We combined scGlycan-seq with scRNA-seq to enable the simultaneous measurement of glycome and transcriptome in single cells, termed scGR-seq (Fig. 1b). Specifically, we employed RamDA-seq, a full-length single-cell total RNA-sequencing method^7^. We performed scGR-seq on the hiPSCs (n = 53) and hiPSC-derived NPCs (11-day differentiation) (n = 43) (Supplementary Tables 6, 7, 8, and 9). After quality control of scRNA-seq data of hiPSCs and NPCs (see Methods and Supplementary Fig. 11), we searched for differentially expressed genes between hiPSC and NPCs. We found that 1,131 and 688 genes were significantly upregulated in hiPSCs and NPCs, respectively (Supplementary Fig. 12, Supplementary Table 10). Consistent with neural differentiation of hiPSCs, GO enrichment analysis demonstrated that gene sets annotated with neuron-associated terms were significantly enriched in NPCs (Supplementary Table 11). Furthermore, transcriptome data of hiPSCs and hiPSC-derived NPCs showed cell-type-specific expression of 41 selected cell-type marker genes (Supplementary Fig. 13)^25,26^. For example, NPCs showed higher expression of neural progenitor marker genes such as *NES* (*NESTIN*), *PAX6*, and *SOX1*^25^ and lower expression of hPSC marker genes such as *NANOG* and *POU5F1^26^*, which agree well with fluorescence staining (Supplementary Fig. 6). These data suggest the scRNA-seq data of GR-seq reflects transcriptome information with biological relevance.

We analyzed the correlation of lectins across cells and found lectins that fluctuated with each other based on their correlation (Supplementary Fig. 14). Lectins were separated into two large clusters: one cluster containing rBC2LCN and TJAII, and another cluster containing other 37 lectins. rBC2LCN and TJAII commonly recognize α1-2fucosylated glycans, which are upregulated in hiPSCs (Supplementary Figs. 10 and 14)^20,27–29^. In contrast, other lectins showed higher binding to NPCs or comparable sigals between the two cell types.

We then addressed how the information from two modalities can be used. When we performed UMAP, a non-linear dimensional clustering, based on only the mRNA or glycan data using the Seurat workflow, the two cell types (hiPSCs and NPCs) were partially separated (Fig. 3a, b). In contrast, when we performed UMAP based on both the mRNA and glycan data using the Seurat weighted nearest neighbor workflow, the two cell types were clearly separated (Fig. 3c). We also found that the unsupervised clustering using both mRNA and glycan data completely agreed with the cell type annotation (Fig. 3f), whereas the clustering based on either mRNA or glycan data showed poorer concordance (Fig. 3d, e). The difference in the clustering results was quantitatively verified by the adjusted Rand index (Fig. 3g). This tendency was also confirmed when we performed PCA based on the mRNA or lectin data and partial least squares (PLS) regression using both mRNA and glycan information, where the latter separated the two cell types clearer than the former two (Supplementary Fig. 15). These results demonstrated how the combination of mRNA and glycan modalities help characterize cell identities.

**Fig. 3:**
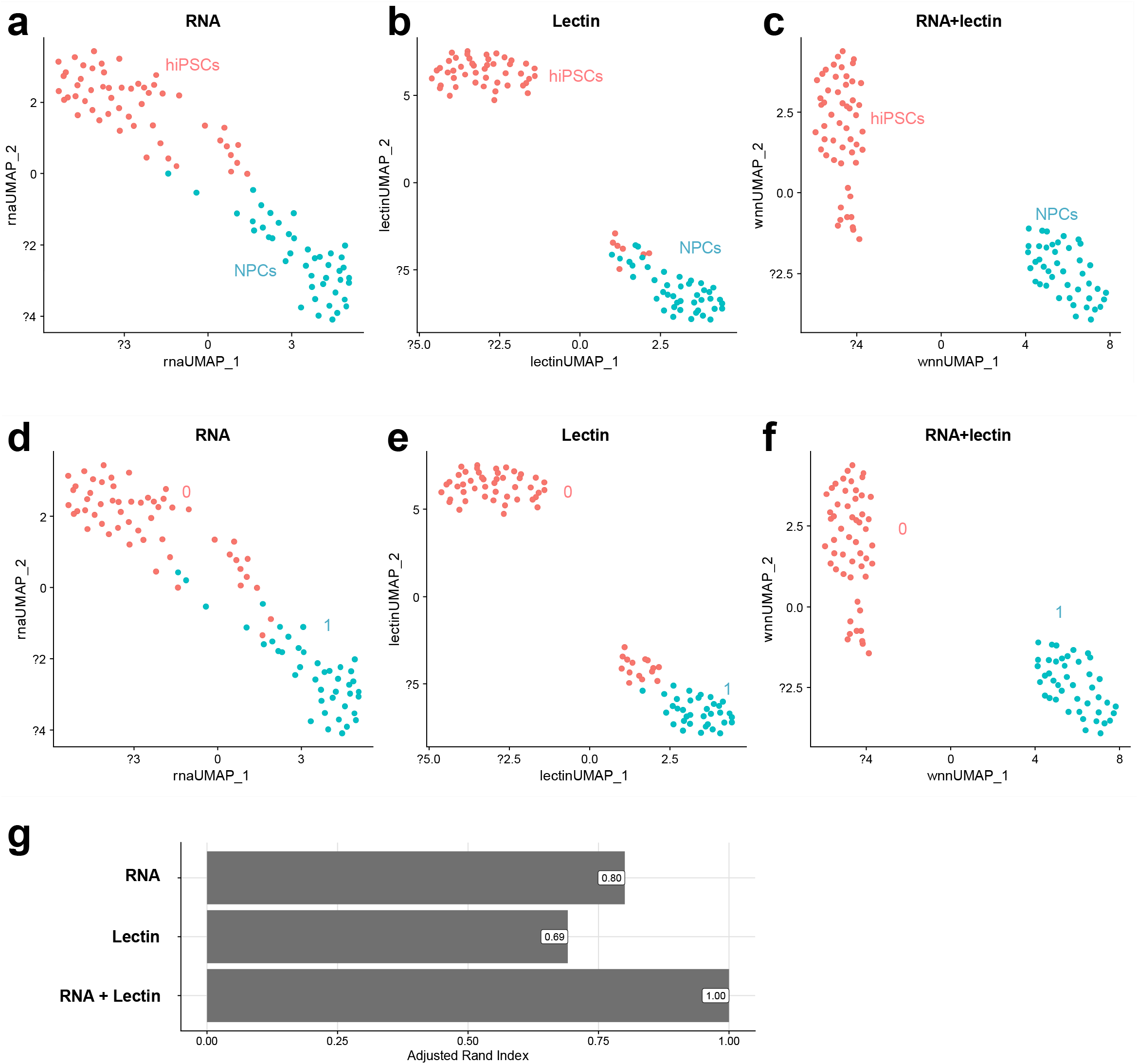
Dimensional reduction and clustering using the Seurat workflow. **(a-c)** UMAP visualization based on (a) only the scRNA-seq data and (b) only the scGlycan-seq, (c) both scRNA-seq and scGlyca-seq (scGR-seq) data of hiPSCs (n = 53, *red*) and NPCs (n = 43, *green*). UMAP computation was performed based on the weighted nearest neighbor graph that integrates the RNA and lectin modalities. **(d-f)** Clustering results based on (d) only scRNA-seq data, (e) only scGlycan-seq data, (f) both scRNA-seq and scGlyca-seq (scGR-seq) data using a shared nearest neighbor (SNN) modularity optimization based clustering algorithm. **(g)** Bar plots showing adjusted Rand index as a measure of the similarity of the clustering in (d), (e), and (f) to the cell type annotation (a-c).

### Relationship between mRNA and glycan in single cells

Simultaneous transcriptome and glycome measurements could associate genes with glycans at single-cell level. The PLS regression analysis described above found a group of mRNAs and lectins that were associated with each other differently per component (see Methods, Fig. 4a). For the component p1, where weights were high for rAAL, rBC2LA, rLSLN, and rBanana, the mRNAs (high p1 weight) related to brain development, cell projection morphogenesis, neural precursor cell proliferation, sensory organ development, and negative regulation of nervous system development were the most enriched gene sets (Fig. 4b, c, Supplementary Table 12), suggesting that the glycan ligands of these lectins might be closely associated with neural differentiation. This analysis allowed us to infer each glycan’s potential functions and roles as a marker through the set of genes associated with the glycan. Furthermore, by summing the loadings of each component, we obtained the overall relationship between lectins and glycosylation-related genes (see Methods and Supplementary Fig. 16). For example, *ST6Gal1* showed the highest correlation with rBC2LCN, consistent with the previous study that both *ST6Gal1* and rBC2LCN ligands are highly expressed in hiPSCs^20^. These exemplify how scGR-seq is useful for finding potential relevance between transcriptome and glycome layers, although further detailed analysis should be performed in future studies.

**Fig. 4:**
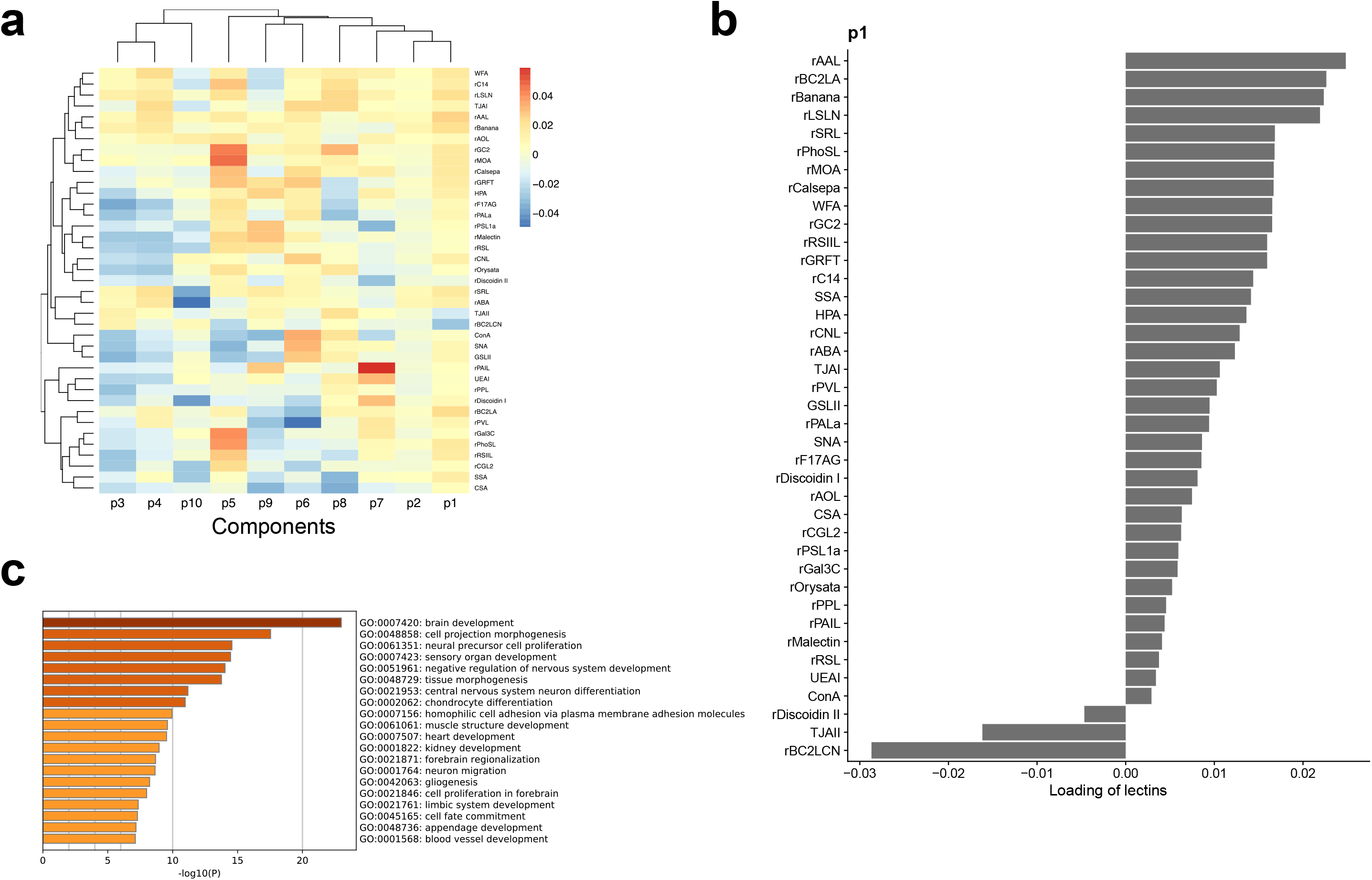
PLS regression. **(a)** A heat map of the association between each lectin and each component inferred by PLS regression. Rows represent lectins and columns represent components. **(b)** Weights of lectins for the component p1 from the PLS regression results. **(c)** Enriched gene sets for the genes associated with the component p1 of the PLS regression results. Gene enrichment analysis was performed with Metascape. The results of gene enrichment analysis are available in Supplementary Table 12.

### scGR-seq reveals pluripotency and differentiation glycan markers

A pseudotime analysis on the mRNA data of hiPSCs and NPCs reconstructed within-cell-type variability reflecting cell differentiation progression (see Methods): expression of hiPSC (*POU5F1*) and NPC (*OTX2*) marker genes showed a progressive decrease and increase along with the pseudotime (generalized additive model, *q* < 0.05) (Fig. 5a), indicating biological relevance of pseudotime reconstruction. Based on the mRNA pseudotime, we next searched for lectins showing changes along with the pseudotime (Fig. 5b; generalized additive model, *q* < 0.05). rBC2LCN, the known pluripotency marker probe^21^, was decreased along pseudotime, while other lectins such as rBanana (mannose binder), which is not known as any cell surface marker, were increased. To confirm the differential binding of rBC2LCN and rBanana, we conducted fluorescence microscopy examination. Consistently, rBC2LCN staining was diminished after differentiation into NPCs (Figs. 2c and 5c). While rBanana showed intracellular organelle staining in hiPSCs, the cell surface was stained in hiPSC-derived NPCs, suggesting a drastic change in the localization of the glycan ligands of rBanana during differentiation (Fig. 5c). Because rBanana showed high weight in component p1 (Fig. 4b) and the component p1 was associated with neuron-related gene sets (Fig. 4c), the glycan ligands of rBanana may be related to neural differentiation.

**Fig. 5:**
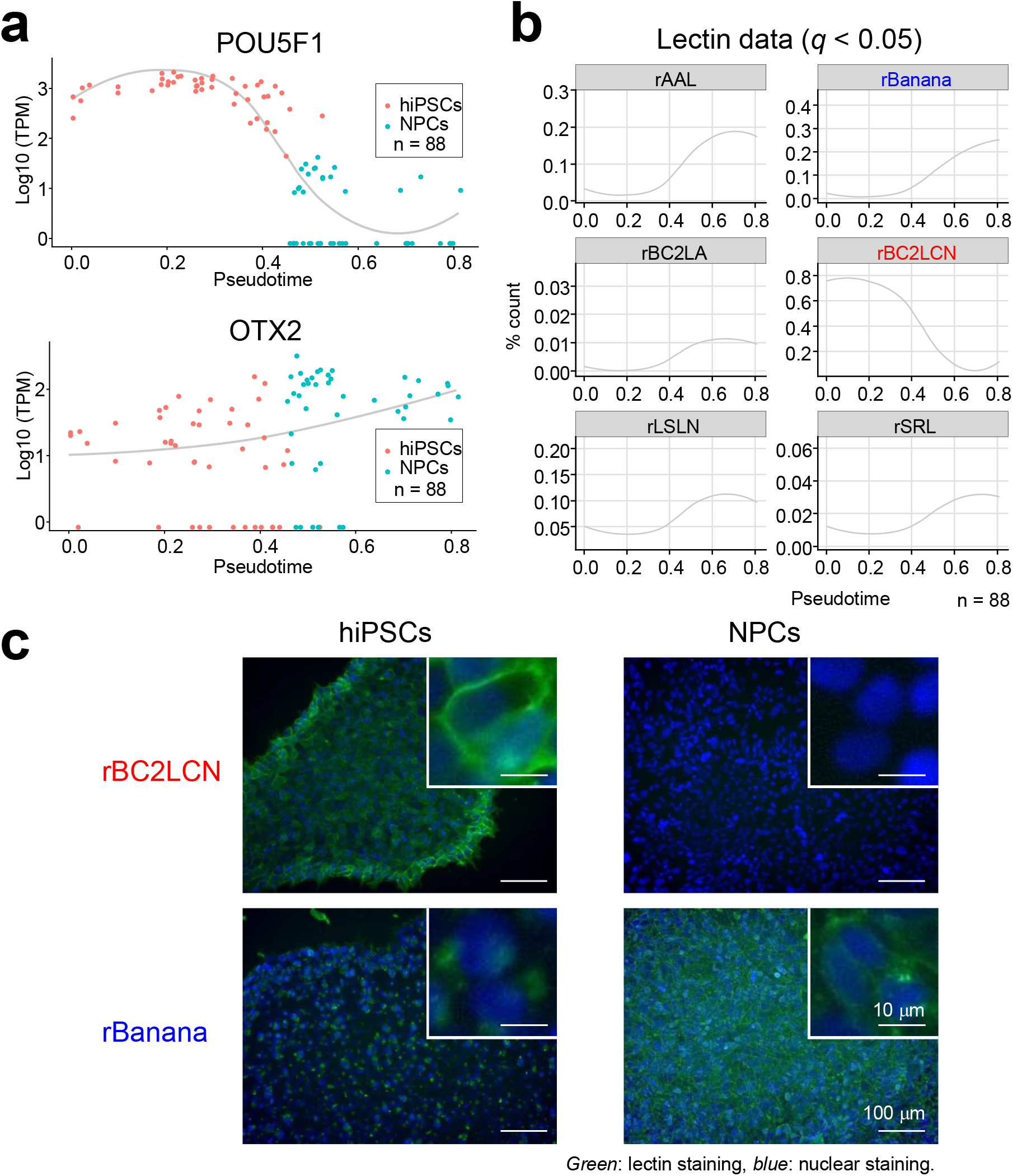
Psuedotime analysis of mRNA and lectin signals. **(a)** Expression profile of the hPSC marker (*POU5F1*) and NPC marker (*OTX2*) genes. The *x*-axis represents the pseudotime inferred by slingshot using mRNA data. The *red* and *blue* points indicate hiPSCs and NPCs, respectively. The curve represents smoothed conditional means calculated by the LOESS (locally estimated scatterplot smoothing) method. **(b)** Dynamically regulated lectins along the pseudotime. The pseudotime is the same as in (a). A generalized additive model was fitted to the lectin signals to search for dynamically regulated lectins (*q* < 0.05 after the multiple testing correction using the Benjamini & Hochberg method). The red and blue points indicate hiPSs and NPCs, respectively. The curve represents smoothed conditional means calculated by the LOESS method. **(c)** Fluorescence staining of hiPSCs and hiPSC-derived NPCs by rBC2LCN and rBanana. hiPSCs and hiPSC-derived NPCs (17-day differentiation) were fixed with 4% paraformaldehyde at room temperature for 20 min. After blocking with PBS containing 1% BSA at room temperature for 30 min, cells were incubated with 1 μg/mL of fluorescently-labeled rBC2LCN and rBanana for 1 h at room temperature and observed by fluorescence microscopy. The nucleus was stained with Hoechst 33342 (x 1,000) at room temperature for 5 min. Insets show high magnification of selected fields. Nucleus: *blue*. Lectin: *green*. Scale bar: 100 μm or 10 μm (insets).

We also examined genes correlated with hiPSC-specific rBC2LCN at the single-cell level (Supplementary Fig. 17 and Supplementary Table 13). The highest positive correlation coefficient (0.77) was observed with the hPSC marker gene (*POU5F1*), whereas the highest negative (−0.65) was observed with the NPC marker gene (*VIM*), supporting the previous finding that rBC2LCN was a maker probe for hPSCs^20,21^.

Furthermore, we searched for lectins correlated with hPSC marker genes (*MYC*, *LEFTY1*, and *LEFTY2*) within hiPSCs (Supplementary Fig. 18), all of which are known to show gene expression variability in pluripotent cells. Expectedly, rBC2LCN exhibited the highest correlation with *MYC*. *LEFTY1* and *LEFTY2* exhibited the highest correlation with rGal3C (galactose binder) and rBC2LA (mannose binder), respectively, suggesting a possible link between glycan ligands of these lectins and the functions of *LEFTY1* and *LEFTY2* in the regulation of self-renewal and differentiation^26,30,31^. These results demonstrate that scGR-seq revealed the coordinated heterogeneity in pluripotency/differentiation markers across mRNA and glycans even within cells of the same cell types.

## Discussion

In this study, we first established glycan profiling by sequencing (Glycan-seq) using 41 DNA-barcoded proteins. Distinct cell populations could be discriminated based on cell surface glycan expression. Relative quantitative differences in expression levels of glycans observed by flow cytometry could be measured by Glycan-seq. No visible non-specific interaction between lectins was observed under the condition used in this study. Furthermore, Glycan-seq was proved to be applicable to single cells. Therefore, Glycan-seq is feasible for profiling cell surface glycans in both bulk and single cells. The amount of DNA barcodes conjugated with lectins affects to the lectin binding signals. However, the aim of the developed method is to compare the lectin binding signals between samples by using the same lot of DNA-barcoded lectin library, which is similar to flow cytometry analysis, other sequencing-based analytical methods such as CITE-seq^16^, and lectin microarray^13^.

We then combined scGlycan-seq with RamDA-seq (full-length total RNA sequencing in single cells), which shows high sensitivity to non-poly(A) RNA and near-complete full-length transcript coverage, to develop scGR-seq, the first method to jointly profile glycan and RNA in single cells. In this study, we focused on the relationship between mRNA expressions and lectin signals and showed that scGR-seq is useful for finding potential relevance between transcriptome and glycome; however, analyzing the relationship between lectin signals with non-poly(A) RNAs, such as nascent RNAs, histone mRNAs, long noncoding RNAs (lncRNAs), circular RNAs (circRNAs), and enhancer RNAs (eRNAs), is also possible and an interesting direction.

In terms of glycan-detection probes, cost-effective and commercially-available lectins, either from natural sources or recombinantly-expressed, were used for Glycan-seq^12,13^. Any glycan-binding probes, such as anti-glycan antibody and glycan-binding peptide, can be incorporated in Glycan-seq^32^.

scGR-seq can be widely adapted to cells, organoids, and tissues and can be applied to various scientific fields such as stem cell biology, immunology, cancer biology, and neuroscience. One of the biggest advantages of scGR-seq is that the method could be applied for the development of novel glycan markers of rare cells such as circulating tumor cells (CTCs), fetal nucleated red blood cells, cancer stem cells.

Identification of glycan markers of rare cells could be performed as follows: (1) rare cells are first isolated and analyzed by scGR-seq. (2) Lectins with specificity to rare cells are selected. Such lectins can be directly used for the identification and concentration of rare cells. (3) Glycoprotein ligands of lectins are identified by LS-MS/MS analysis of lectin-precipitated fractions. (4) Monoclonal antibodies recognizing both a glycan epitope and a peptide sequence could be generated as specific probes for rare cells as previously reported^33^. Such antibodies would be useful for the development of novel diagnosis^34^ and antibody drugs^35^. scGR-seq can also be applied to cells derived from other organisms such as fungi and bacteria using the available platform, as lectins can bind glycans expressed by the cells of any organism^36,37^.

There are limitations in scGlycan-seq and scGR-seq. Similar to flow cytometry and lectin microarray, absolute amounts of glycans and accurate glycan structures cannot be determined directly from the signal intensities as described above. Another limitation of the current system is the throughput. In this sense, scGR-seq can be improved by the adaption to droplet^38^, nano-well^39^, and indexing-based high-throughput single-cell technologies^40^ in future studies. We believe that it has the potential to advance the understanding of cellular heterogeneity and the biological roles of glycans across diverse multicellular systems across species.

## Methods

### Cells

201B7 hiPSCs were cultured in 2 ml of mTeSR Plus (VERITAS, Tokyo, Japan) on a 6-well plate coated with Matrigel (Corning International, Tokyo, Japan)^26^. 201B7 hiPSCs were differentiated to neural progenitor cells using STEMdiff SMADi Neural Induction Kit (VERITAS).

### Conjugation of lectins to DNA oligonucleotides

100 μg of lectin in 100 μl of PBS was incubated with ten-times the molar amount of dibenzocyclooctyne-N-hydroxysuccinimidyl ester (DBCO-NHS) (Funakoshi Co., Ltd., Tokyo, Japan) at 20°C for 1 h under dark (Supplementary Fig. 1). 10 μl of 1 M Tris was then added and incubated at 20°C for 15 min under dark to inactivate the excess DBCO-NHS. The excess DBCO-NHS was also removed by using Sephadex G-25 desalting columns (GE Healthcare Japan Co., Tokyo, Japan). The resulting DBCO-labeled lectin was then incubated at 4 °C with ten-times the molar amount of 5’-azide-modified DNA oligonucleotides (Integrated DNA Technologies, KK, Tokyo, Japan). To remove unbound oligonucleotides and obtain lectins with sugar-binding activity, the DNA-barcoded lectin was purified by affinity chromatography using appropriate sugar-immobilized Sepharose 4B-CL (GE) based on the glycan-binding specificity of each lectin. Finally, DNA-barcoded lectin was dialyzed by Tube-O-DIALYZER (Takara Bio Inc., Shiga, Japan) and concentrated using centrifugal filters having a 10 kDa molecular weight cut off (MWCO) (Merck KGaA, Darmstadt, Germany). The purified DNA-barcoded lectin was analyzed by SDS-PAGE. Protein and DNA concentration were measured using the Bradford protein assay (Bio-Rad Laboratories, Inc., CA, USA) and Quant-iT OliGreen ssDNA Reagent (Thermo Fisher Scientific KK, Tokyo, Japan).

### List of lectins

See Supplementary Table1 for a list of lectins and barcodes used for bulk and single cell Glycan-seq and GR-seq.

### Glycan-seq in bulk and single cells

Cells (1×10^5^) were suspended in phosphate-buffered saline (PBS) containing 1% bovine serum albumin (BSA) and incubated with 41 DNA-barcoded proteins at a final concentration of 0.5 μg ml^-1^ at 4°C for 1 h. After washing three times with 1 ml of PBS/BSA, cell number was counted by TC20 auto cell counter (Bio-Rad Laboratories, Inc.). Cells (1×10^4^ cells per tube or 1 cell per tube) were distributed into a PCR tube (FCR&BIO Co., LTD., Hyogo, Japan). One cell dispense was performed using TOPick I Live Cell Pick system (Yodaka Giken, Kanagawa, Japan). To liberate oligonucleotides from cells, the cells were irradiated at 365 nm, 15 W for 15 min using UVP Blak-Ray XX-15L UV Bench Lamp (Analytik Jena, Kanagawa, Japan).

The liberated oligonucleotides were then amplified using NEBNext Ultrall Q5 (New England BioLabs Japan Inc., Tokyo, Japan), and i5-index and i7-index primers containing cell barcode sequences (Supplementary Table 14). PCR reactions were performed as follows: denaturing (45 s at 98°C, 1 cycle), amplification (10 s at 98°C, 50 s at 65°C, 20 cycles), extension (5 min at 65°C, 1 cycle). The PCR products were then purified by the Agencourt AMPure XP kit (Beckman Coulter, Inc., Tokyo, Japan), followed by the manufacturer’s protocol. The size and the quantity of the PCR products were analyzed by MultiNA (Shimadzu Co., Kyoto, Japan). The PCR products (4 nM from every single cell) were treated with the Miseq Reagent Kit v2 50 Cycles (Illumina KK, Tokyo, Japan) and sequenced by the MiSeq sequencer (26 bp, paired-end) (Illumina KK).

### Data analysis of Glycan-seq

DNA barcodes derived from lectins were directly extracted from the reads in the FASTQ format. The number of DNA barcodes bound to each cell was counted using the developed software, Barcode DNA counting system (Mizuho Information & Research Institute, Inc., Tokyo, Japan). In order to match the sequence, the first three bases were removed. As maximum, two mismatches in flanking region and one mismatch in middle region were accommodated. Each DNA barcode count was divided by the total lectin barcode count and expressed as a percentage (%) for each lectin. In order to filter out the cells with low total barcode count, the cut-off value of total lectin barcode count was determined using Otsu’s method^41^. The cut-off value was determined for each cell type, and hFibs = 19,465, hiPSCs = 6,126, Day4 = 14,723 and Day11 = 8,798 were used for the analysis (Fig. 2). No bias for Principal component analysis (PCA) due to the total barcode count was observed for scGR-seq data of hiPSCs and NPCs. PCA was performed to simplify the multivariate data of glycan-seq by reducing the dimensionality. PCA was carried out by SPSS Statistics 19 (IBM Japan, Tokyo, Japan). Data were preprocessed using mean-centering prior to this analysis. Using the first two principal component (PCs), each sample data was visualized on two dimensional plot.

To test the difference of the rBC2LCN signal in scGlycan-seq between hiPSCs and hFibs, and hiPSCs and NPCs, we used the Brunner-Munzel test, an independent two-sample test that assumes neither normality nor homoscedasticity^41^, using the ‘brunner.munzel.test’ function in ‘lawstat’ R package.

### scRNA-seq

Single-cell cDNA library was prepared using the GenNext RamDA-seq Single Cell Kit (Toyobo Co., LTD. Tokyo, Japan), followed by the manufacturer’s protocol^42^. The obtained library was then quantified by real-time PCR using the PowerUp SYBR Green Master Mix (Thermo Fisher Scientific KK) and CFX Connect System (Bio-Rad Laboratories, Inc.). DNA standards for library quantification (Takara Bio Inc., Shiga, Japan) were used as the standard. 10 nM of the cDNA library from every single cell was treated with the NovaSeq 6000 S4 reagent kit and sequenced by Nova-Seq 6000 (151 bp, paired-end) (Illumina KK).

### Preprocess of scRNA-seq data

The FASTQ quality check of reads was performed using FastQC (version 0.11.5). The reads were trimmed by using the fastq-mcf (version 1.0) with the parameters “-L 150 -l 50 -k 4.0 -q 30”. The trimmed reads were mapped to the human reference genome (GRCh38, primary assembly) using HISAT2 (version 2.1.0) with the parameters “--no-softclip”^43^. The resulting SAM files were then converted into BAM files using SAMtools (version 1.4). RSeQC (version 1.2) was used for the quality check of the read-mapping^44^. To mitigate the difference in the number of mapped reads among single cells, the reads in each BAM file were subsampled to 555,421 reads when the BAM file contained more than 555,421 reads using samtools view ‘-s’ option. The featureCounts command in Subread (version 1.6.4) was used to quantify gene-level expression with the parameters “-M -O --fraction -p” and the reference gene model (GENCODE v32, primary assembly)^45^. Transcript per million (TPM) values were calculated from the gene-level expression quantification result using R (version 3.6.1)^46^. R was used for the statistical analyses and figure generation.

### Pseudotime analysis

The cells with low mapped reads were discarded from the pseudotime analysis. Diffusion maps were performed using the destiny package (version 3.0.1) after retaining genes where at least 3 counts were found in at least 10 cells^47^. The pseudotime analysis was performed using the slingshot package (version 1.4.0)^48^ as follows: (1) we selected expressed genes with a count of at least 3 in at least 10 cells. (2) We normalized the count matrix of the expressed genes with the full quantile normalization. (3) We performed diffusion maps on the log-transformed normalized count matrix with the pseudocount of 1 using the “destiny” R package (version 3.0.1). (4) We applied the slingshot analysis on the result of diffusion maps (diffusion components 1 and 2) to estimate the pseudotime. The package gam (version 1.16.1) was used for fitting a generalized additive model with the parameter “family☐=☐Gaussian(link☐=☐identity)” to search for dynamically regulated genes (from highly variable genes (FDR < 0.01)) and lectins (*q* < 0.05 after the multiple testing correction using the Benjamini & Hochberg method).

### Seurat analysis

The Seurat package (version_3.9.9.9014) was used to analyze scGR-seq data for hiPSCs and NPCs. The TPM matrix for the scRNA-seq data and the lectin expression matrix for the scGlycan-seq data was used as inputs. The RNA data was preprocessed using the ‘NormalizeData’ function with the default parameters, the ‘FindVariableFeatures’ function with the parameter ‘selection.method = “vst”’, and the ‘ScaleData’ with the default parameters. The lectin data was preprocessed using the ‘NormalizeData’ function with the parameters ‘normalization.method = ‘CLR’, margin = 2’, the ‘FindVariableFeatures’ function with the default parameters, and the ‘ScaleData’ with the default parameters. Principal component analysis (PCA) was then performed on the RNA data and the lectin data separately using the ‘RunPCA’ function with the parameter ‘approx=FALSE’. Then, UMAP visualization was computed based on the PCA result of RNA (PC1 to 10) and lectin (PC1 to 20) data individually using the ‘RunUMAP’ function with the default parameters. In parallel, the ‘FindNeighbors’ function with the default parameters was used on the PCA result of RNA (PC1 to 10) and lectin (PC1 to 20) data to define *k* nearest neighbor cells with the parameter ‘resolution = 0.5’. To search differentially expressed genes, ‘FindMarkers’ function with the parameter ‘logfc.threshold=0.25’ was performed and genes which match the criteria, Benjamini-Hochberg adjusted p-value < 0.05, were visualized by ‘DoHeatmap’ function. GO enrichment analysis was performed using the DAVID Bioinformatics Resources 6.8 (https://david.ncifcrf.gov/tools.jsp).

To jointly analyze the RNA and lectin modalities, the weighted nearest neighbor (WNN) workflow in Seurat was performed^49^. First, to learn cell-specific modality ‘weights’ and construct a WNN graph that integrates the two modalities, the ‘FindMultiModalNeighbors’ function was used on the PCA results of mRNA (PC1 to 10) and lectin (PC1 to 20) with the parameters ‘knn.range = 50, k.nn=10’. Then, UMAP visualization was computed based on a weighted combination of RNA and lectin data using the ‘RunUMAP’ function with the default parameters. In parallel, graph-based clustering was performed based on the WNN graph using the ‘FindClusters’ function with the parameter ‘resolution = 0.2’. To evaluate the similarity of the clustering with the cell type annotation, the adjusted Rand index was calculated using the ‘mclust’ R package (version 5.4.6).

### PLS regression of scRNA-seq and scGlycan-seq data

The package ropls (version 1.18.8) was used for PLS regression with scRNA-seq and scGlycan-seq data ^50^. Let *X* be *n* × *m* gene expression matrix (scRNA-seq) and Y be *n* × *p* lectin matrix (scGlycan-seq), where *n* is the number of cells, *m* is the number of genes, and p is the number of lectins. PLS regression decomposes *X* and *Y* to maximize the covariance between *T* and *U* as follows:

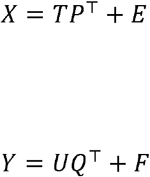

where *T* and *U* are *n* × *l* matrices which is the projection of *X* and *Y*, respectively, and *P* and *Q* are *m* × *l* and *p* × *l* orthogonal loading matrices, respectively, and matrices *E* and *F* are the error terms. The *l* components are analogous to principal components. The associations of genes and lectins were calculated as *PQ^T^* (Fig. 5). The glycogene annotation was retrieved from GlycoGene Database (GGDB, https://acgg.asia/ggdb2/)^51^.

### Flow cytometry

Lectins were recombinantly expressed in *Escherichia coli* and purified by affinity chromatography as described^20^. Lectins were labeled with R-phycoerythrin (PE) using the R-phycoerythrin Labeling Kit (Dojindo Laboratories Co. Ltd., Kumamoto, Japan). 1×10^5^ cells were suspended in 100 μl of PBS/BSA and incubated with 10 μg ml^-1^ of PE-labeled lectins on ice for 1 h. For intracellular staining, cells were fixed with 4% paraformaldehyde at room temperature for 10 min and permeabilized with 0.1% saponin in PBS at room temperature for 10 min. Cells were then incubated with anti-PAX6 mAb (x20, clone No. O18-1330, BD Biosciences, CA, USA) on ice for 1 h. Flow cytometry data were acquired on a CytoFLEX (Beckman Coulter, Inc., CA) and analyzed using the FlowJo software (FlowJo, LLC., OR).

### Fluorescence staining

Cells were fixed with 4% paraformaldehyde at room temperature for 20 min. After blocking with PBS containing 1% BSA and 0.2% Triton-X at room temperature for 30 min, the cells were stained with anti-Nestin mAb (71.1 μg/ml, Clone 25/NESTIN (RUO), BD Biosciences), anti-PAX6 mAb (82.1 μg/ml, clone No. O18-1330, BD Biosciences) or 1 μg/mL of Cy3-labeled lectins at room temperature for 60–90 min. Thereafter, the cells were observed by fluorescence microscopy (IX51, Olympus, Tokyo, Japan). The nucleus was stained with Hoechst 33342 (x 1,000, Dojindo Laboratories).

### Quantitative RT-PCR (qRT-PCR)

Total RNA was extracted using RNeasy Mini Kit (QIAGEN, Hilden, Germany) according to the manufacturer’s protocol. Equal amounts of total RNA from each samples were then reverse-transcribed into cDNA using QuantiTect Reverse Transcription Kit (QIAGEN). qRT-PCR was performed using PowerUp SYBR Green Master Mix (Thermo Fisher Scientific) and CFX Connect (Bio-Rad Laboratories, Inc.). The mRNA expressions of indicated genes were normalized to that of the GAPDH gene. The following primers were used for qRT-PCR: GAPDH-fw(forward): GGCTGGCATTGCCCTCAACG; GAPDH-rv(reverse): AGGGACTCCCCAGCAGTGAG; OCT4-fw: GGAAGGAATTGGGAACACAAAGG; OCT4-rv: AACTTCACCTTCCCTCCAACCA; SOX1-fw: GTCCATCTTTGCTTGGGAAA; SOX1-rv: TAGCCAGGTTGCGAAGAACT; NESTIN-fw: CAGCGTTGGAACAGAGGTTGG; NESTIN-rv: TGGCACAGGTGTCTCAAGGGTAG; PAX6-fw: GTCCATCTTTGCTTGGGAAA; PAX6-rv: TAGCCAGGTTGCGAAGAACT; FOXG1-fw: GCGCAAATGCCGCATAAAT; FOXG1-rv: AAACACGGGCATATGACCACAG.

## Acknowledgments

This work was supported by PRESTO from Grant Number JPMJPR16F6 from Japan Science and Technology Agency (JST), and Adaptable and Seamless Technology transfer Program through Target-driven R&D (A-STEP) Grant Number JPMJTR20UD from JST awarded for HT, and JSPS KAKENHI (Grant Number: 19K20394) and JST-MIRAI Program (Grant Number: JPMJMI18G4) awarded for HO. We thank Tadashi Kimura in National Institute of Advanced Industrial Science and Technology (AIST) for NGS analysis, Keiko Hiemori, Kayo Kiyoi, and Jinko Murakami in AIST for technical assistance on the single cell analysis. We also thank Mika Yoshimura, Tetsutaro Hayashi, and Itoshi Nikaido in Laboratory for Bioinformatics Research, RIKEN Center for Biosystems Dynamics for thoughtful discussion on RamDA-seq data analysis.

## Author contributions

HT designed research, analyzed the data, and wrote the paper. FM carried out the experiments and analyzed the data. HO and HO carries out the experiments, performed data analysis, and wrote the paper.

## Data availability

All raw data of Glycan-seq are provided in the supplementary table. All raw data of scRNA-seq have been deposited to the gene expression omnibus (GEO) with the accession code GSE151642.

