## Supplementary Figure for "Integrated analysis of glycan and RNA in single cells"

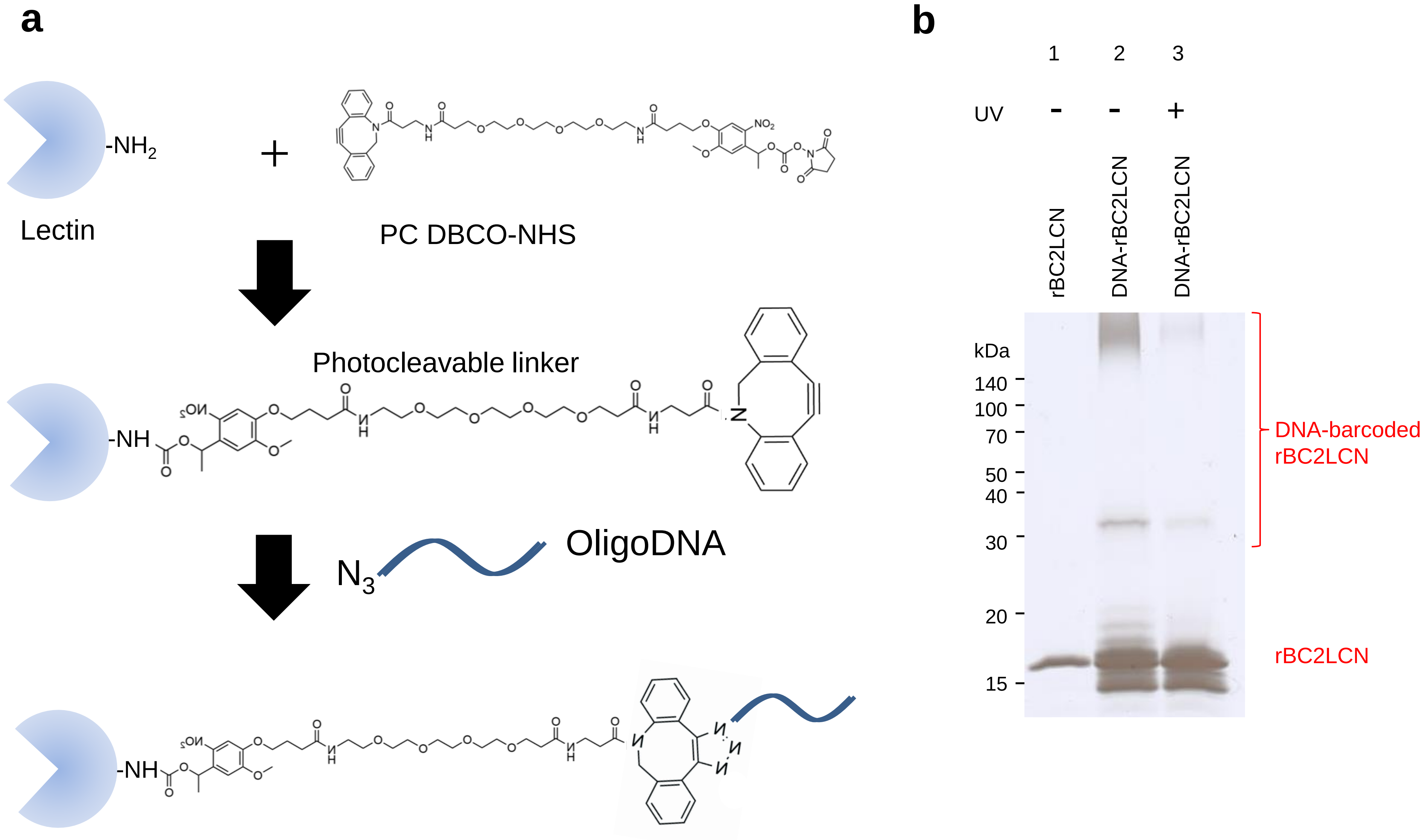


**Supplementary Fig.1 Conjugation of lectins to DNA oligonucleotides.**
(a) Illustration of the protocol of the conjugation of lectins with DNA oligonucleotides. (b) rBC2LCN shows a single band at 16 kDa (lane 1). DNA-barcoded rBC2LCN exhibited a high-molecular weight smear band at >140 kDa (lane 2). Cleavage of DNA barcodes from rBC2LCN by UV exposure collapses the smear to the MW of rBC2LCN (16 kDa) (lane 3).

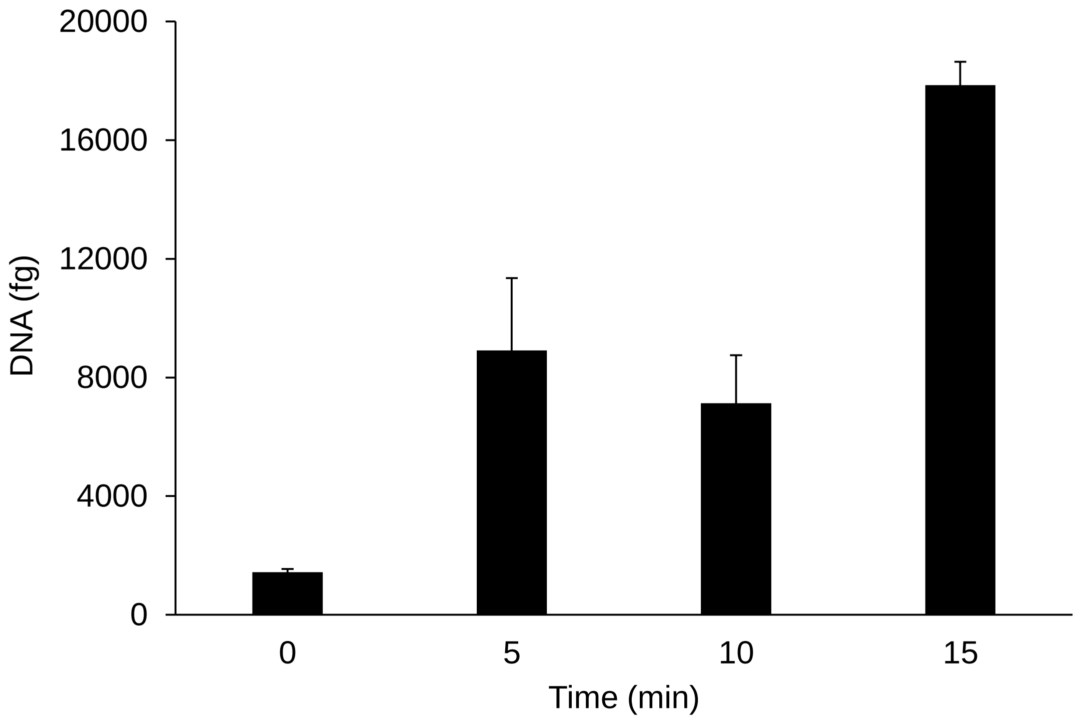


**Supplementary Fig. 2 Optimization of UV exposure time to release DNA barcode from DNA-barcoded lectins.**

Capan-1 was incubated with 1 μg/mL of DNA-barcoded rBC2LCN on ice for 1 h. UV light was exposed to cells for 0, 5, 10, 15 min and the released DNA was recovered and analyzed by qRT-PCR.

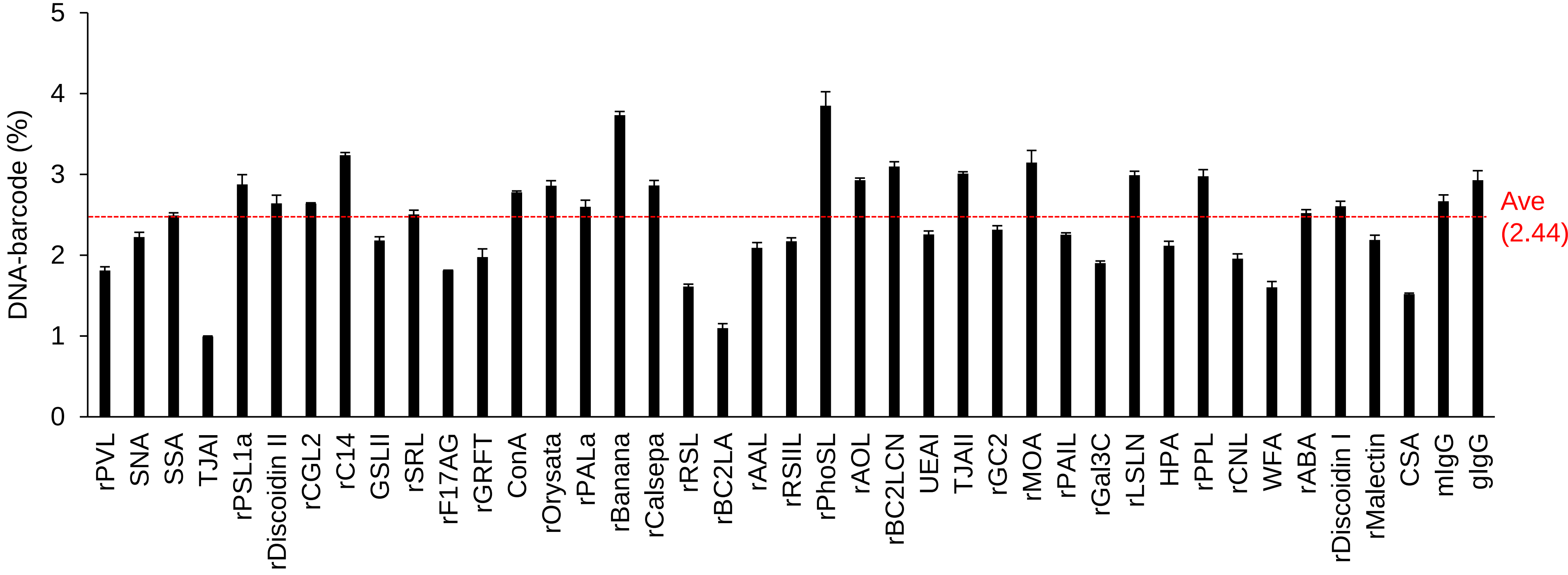


**Supplementary Fig. 3 PCR amplification bias of DNA barcodes.**
DNA barcode (1 pM) of 41 probes (lectins and antibodies) was mixed and amplified using NEBNext UltraII, and i5-index and i7-index primers. The PCR products were then purified by the Agencourt AMPure XP kit, followed by the manufacturer's protocol. The size and the quantity of the PCR products were analyzed by MultiNA. The PCR products (4 nM for each DNA barcode) were treated with the Miseq Reagent Kit v2 50 Cycles and sequenced by the MiSeq sequencer (26 bp, paired-end). DNA barcodes were directly extracted from the reads in the FASTQ format. The number of DNA barcodes was counted using the developed software, Barcode DNA counting system. Each DNA barcode count was divided by the total lectin barcode count and expressed as a percentage (%) for each lectin.

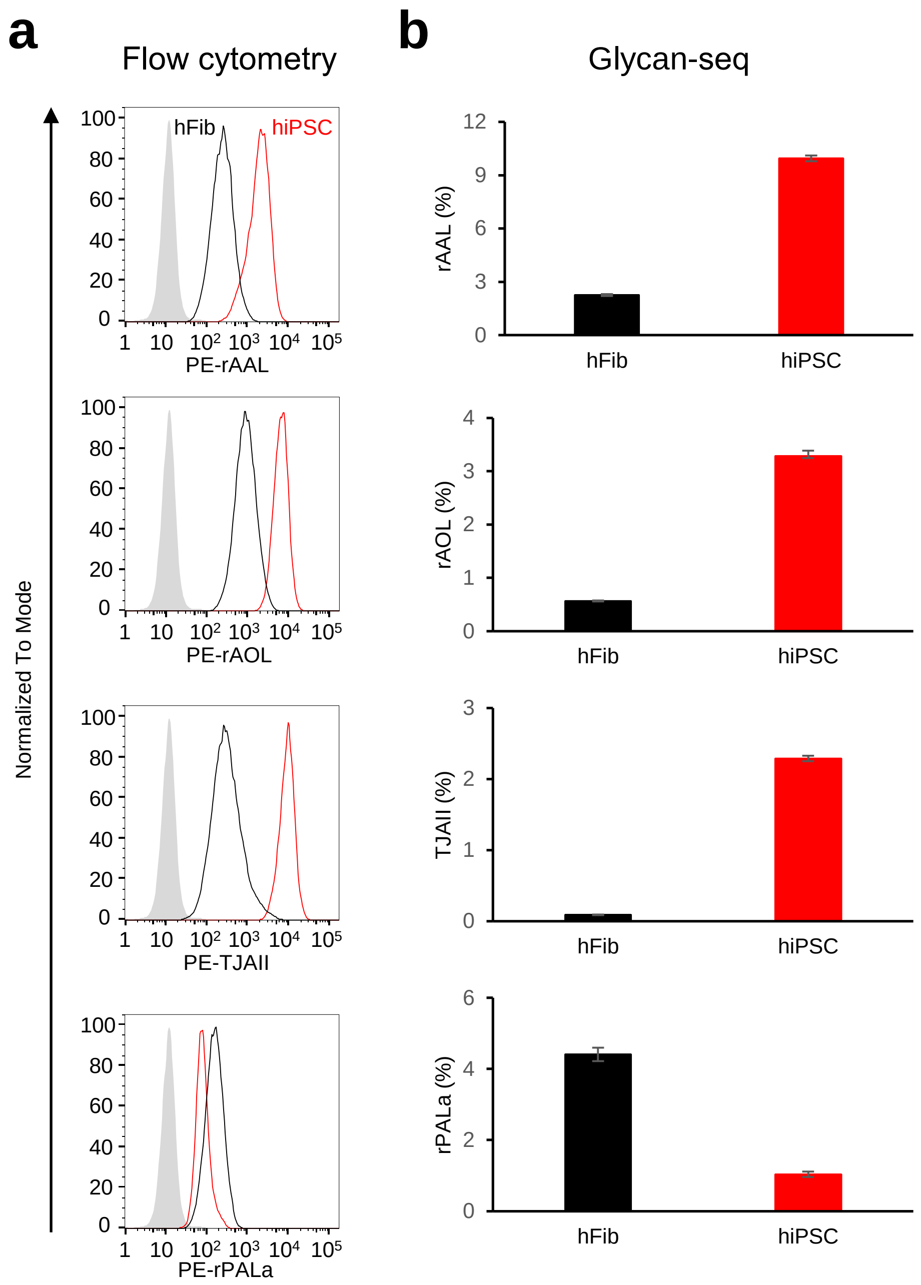


**Supplementary Fig.4 Comparison of bulk Glycan-seq and flow cytometry.**

(a) Binding of R-phycoerythrin (PE)-labeled lectins to hiPSCs (*red line*) and hFibs (*black line*) was analyzed by flow cytometry. *Grey*: Binding of PE-labeled BSA to hiPSCs (negative control). (b) Binding of DNA-barcoded lectins to hiPSCs and hFibs was analyzed by Glycan-seq. The number of DNA-barcode derived from rBC2LCN was divided by that of the DNA-barcode of all lectins, multiplied by 100, and expressed as percentage (%). Data are shown as average ± SD of triplicates of the same sample.

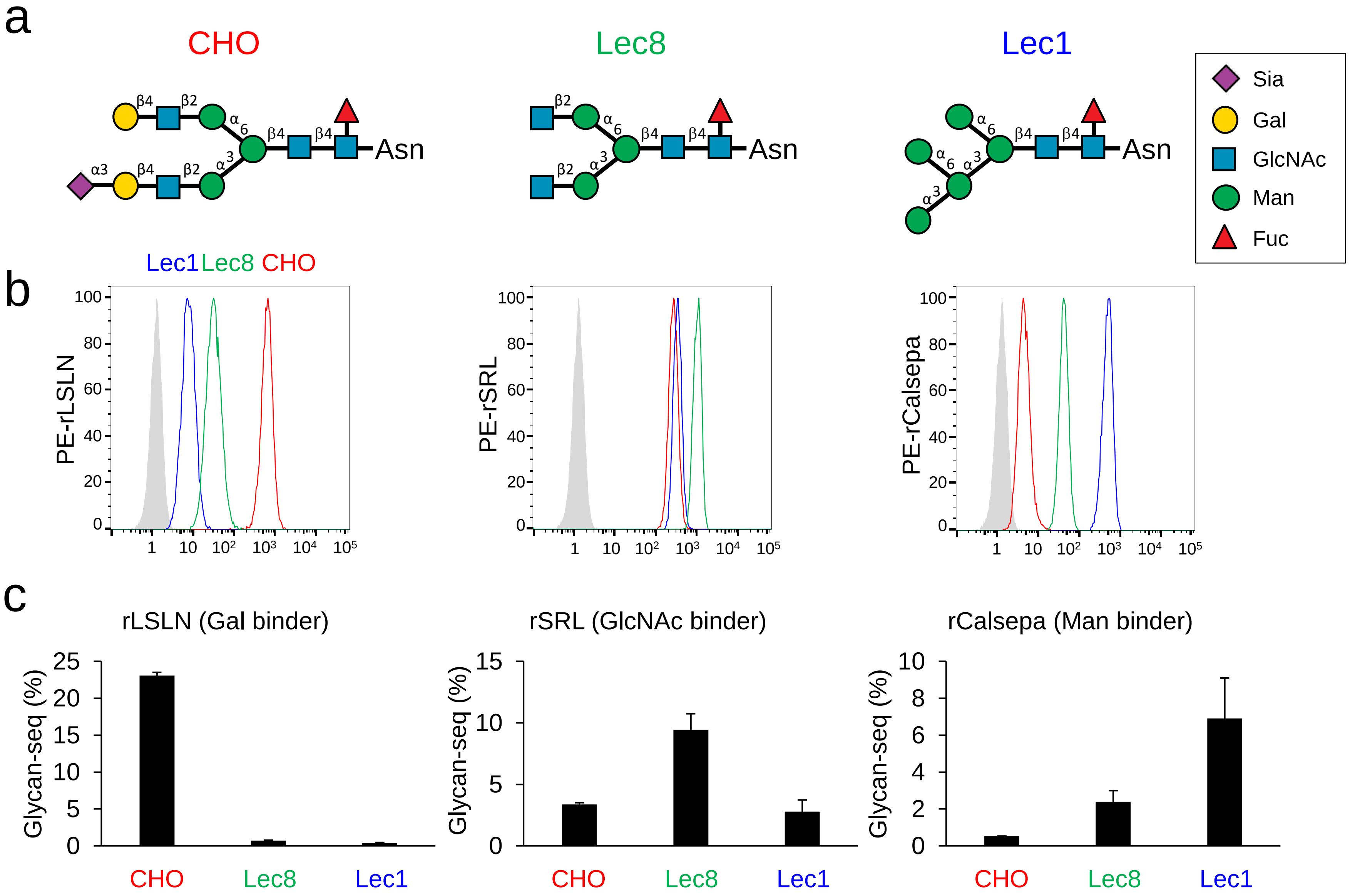


**Supplementary Fig.5 Glycan profiling of CHO, Lec8, and Lec1 by Glycan-seq.**
(a) Typical N-glycan structures expressed in CHO, Lec8, and Lec1. (b) Binding of Gal-binder (rLSLN), GlcNAc-binder (rSRL), and Man-binder (rCalsepa) to CHO, Lec8, and Lec1 analyzed by flow cytometry. (c) Binding of Gal-binder (rLSLN), GlcNAc-binder (rSRL), and Man-binder (rCalsepa) to CHO, Lec8, and Lec1 analyzed by GR-seq. Data are shown as average ± SD of triplicates of the same sample.

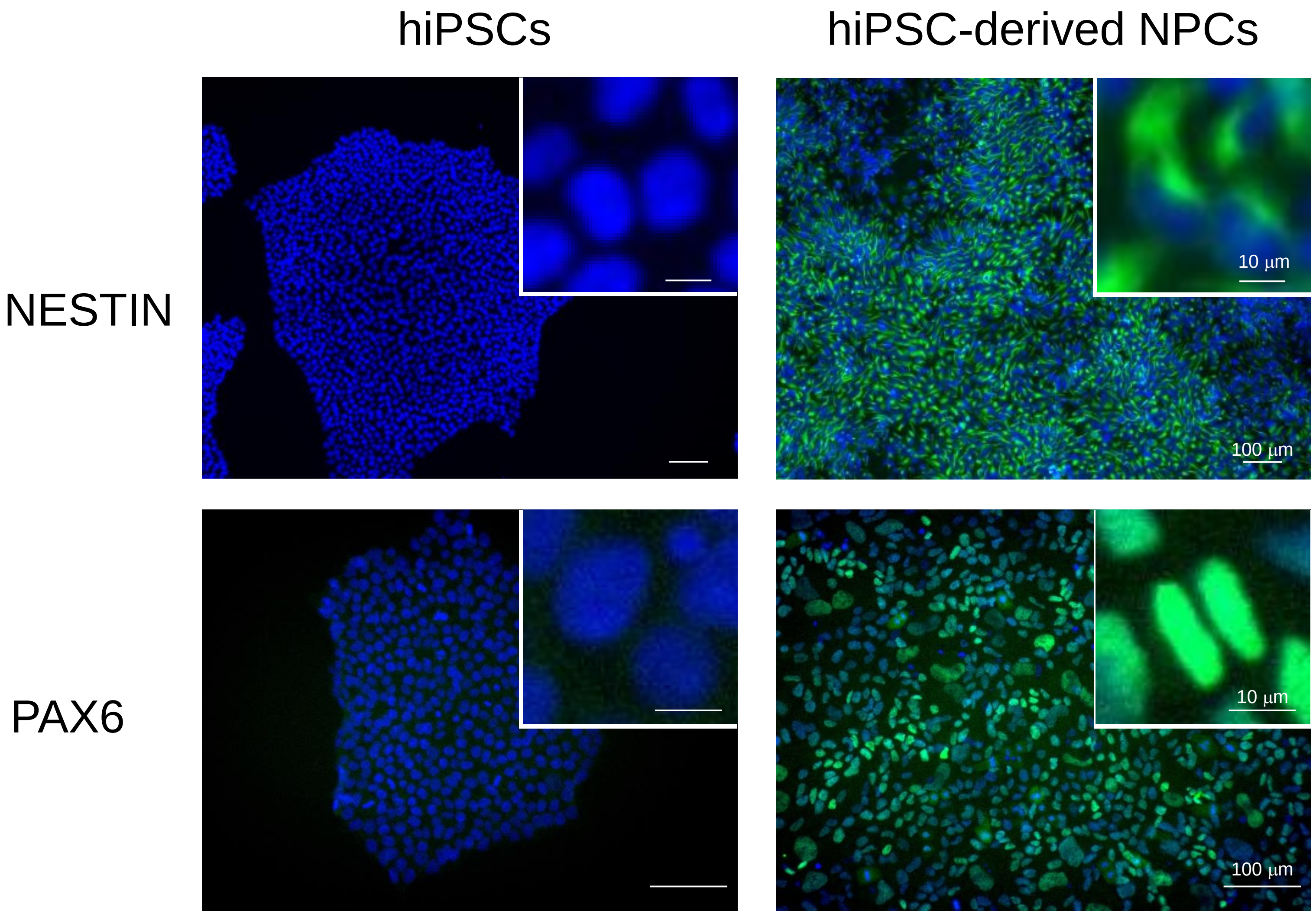


*Green*: antibody staining. *Blue*: nuclear staining.

**Supplementary Fig.6 Fluorescence staining of hiPSCs and hiPSCs-derived NPCs by anti-NESTIN or PAX6 mAb.** hiPSCs and hiPSC-derived hNPCs (Nestin: 17-day differentiation, PAX6: 25-day differentiation) were fixed with 4% paraformaldehyde at room temperature for 20 min. After blocking with PBS containing 1%BSA and 0.2% Triton-X at room temperature for 30 min, cells were stained with anti-Nestin mAb (71.1 μg/ml) or anti-PAX6 mAb (82.1 μg/ml) and Hoechst33342 (1 μg/ml) at room temperature for 90 min. Insets show high magnification of selected fields. Nuclear: *blue*. Nestin and PAX6: *green*. Scale bar: 100 μm or 10 μm (Insets).

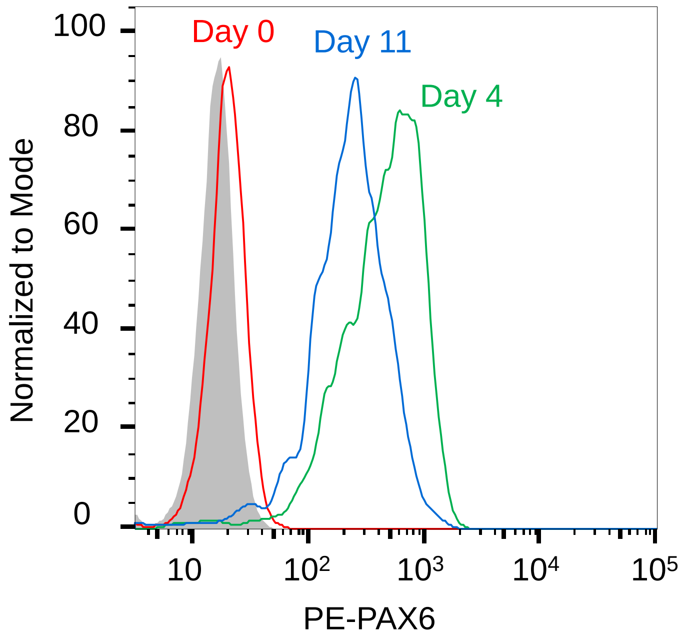


**Supplementary Fig.7  Flow cytometry analysis of hiPSCs during differentiation into NPCs**

hiPSCs before (Day 0, *red line*) and after differentiation into neural progenitor cells (Day 4, *green line*; Day 11, *blue line*) were fixed with 4% paraformaldehyde at room temperature for 10 min and permeabilized with PBS containing 0.1% saponin at room temperature for 10 min. Cells were stained with PE-labeled anti-PAX6 mAb (clone No. O18-1330) on ice for 1 h and analyzed by flow cytometry.

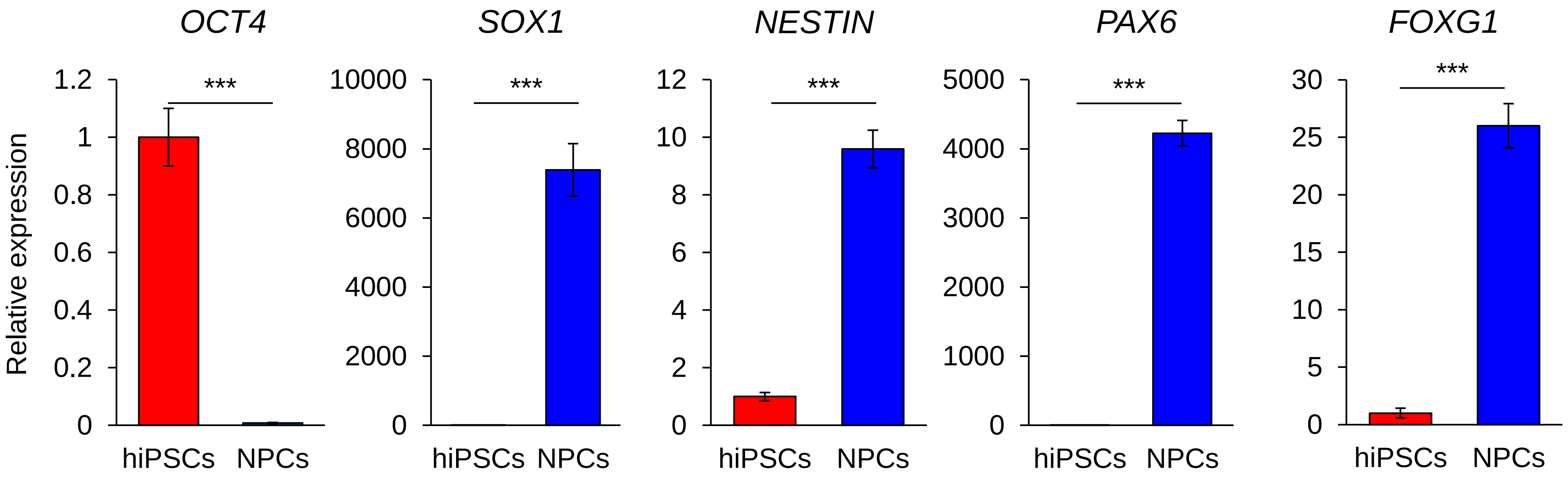


**Supplementary Fig.8 Relative expression of mRNA of hPSC (OCT4) and NPC markers (SOX1, NESTIN, PAX6, FOXG1) in hiPSCs and hiPSC-derived hNPCs.**
The expression of mRNA was measured by qRT-PCR. Data are shown as relative to hiPSC data. Data are shown as average ± SD of triplicate experiments. *** $p < 0.001$. *t-test.*

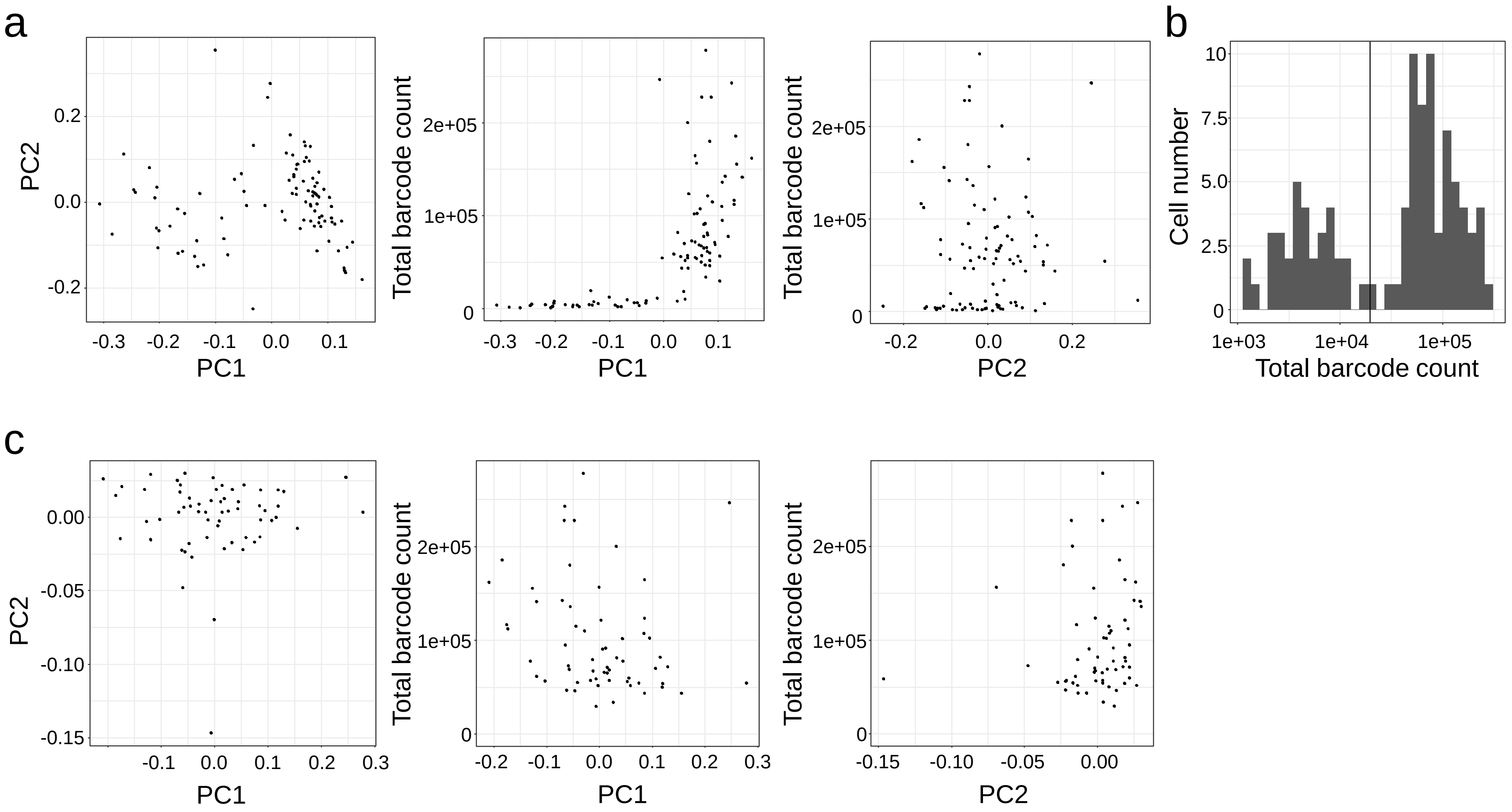


**Supplementary Fig. 9 Quality control of scGlycan-seq data.** (a) PCA plot of scGlycan-seq data of fibroblasts (n = 96) (*left*). The relationship of PC1 or PC2 with total barcode counts was analyzed (*middle*, *right*). (b) Histogram of total barcode count of fibroblasts. Black line indicates threshold determined by Otshu's method (barcode count > 19465). (c) PCA plot of scGlycan-seq data of fibroblasts after removal of cells with low total barcode counts (*left*). The number of samples was reduced from 96 to 61. The relationship of PC1 or PC2 with total barcode counts was illustrated (*middle*, *right*). No obvious bias was observed in these data sets.

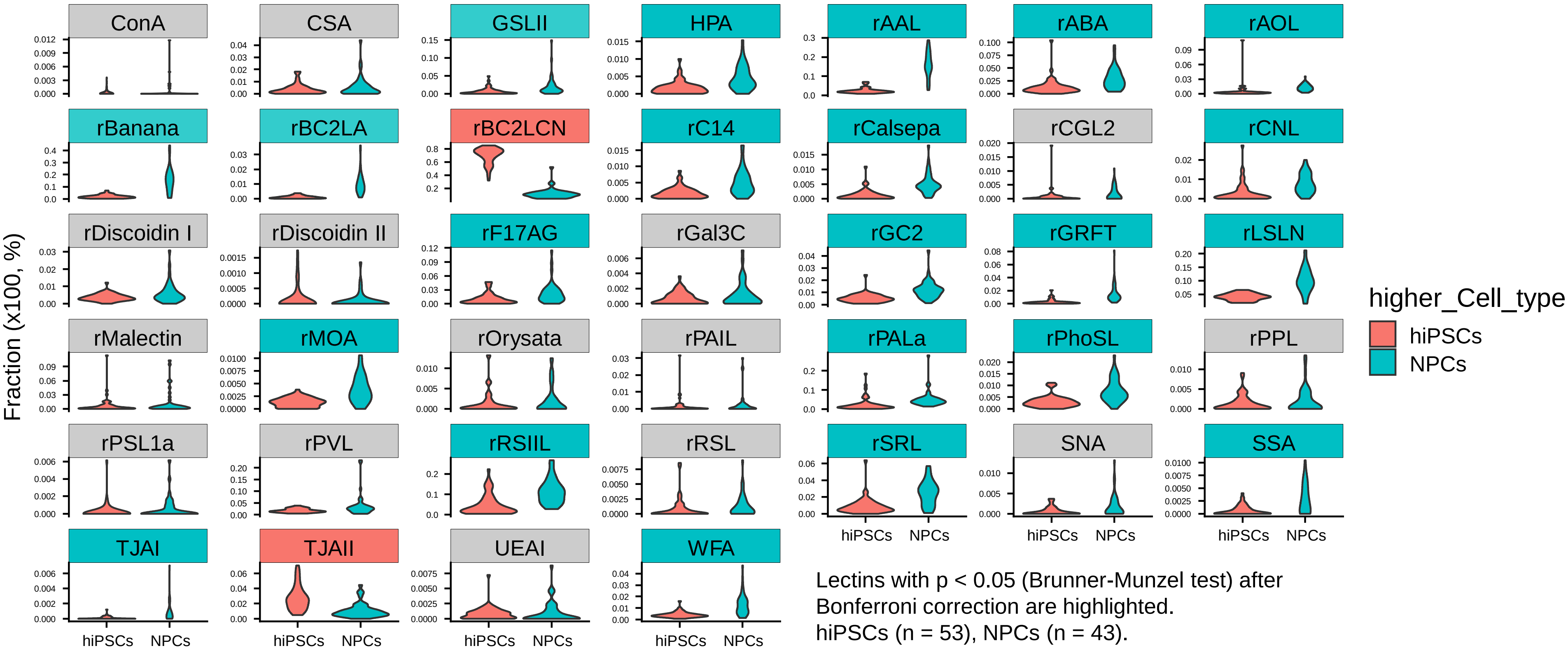


**Supplementary Fig.10 scGlycan-seq data of hiPSCs and NPCs.**
Violin plots showing the signal level of each lectin in scGlycan-seq data of hiPSCs (n = 53), NPCs (n = 43). Lectins with p < 0.05 (Bunner-Munzel test) after Bonferoni correction are highlighted in red and cyan when the average signal was higher in hiPCSs and NPCs, respectively.

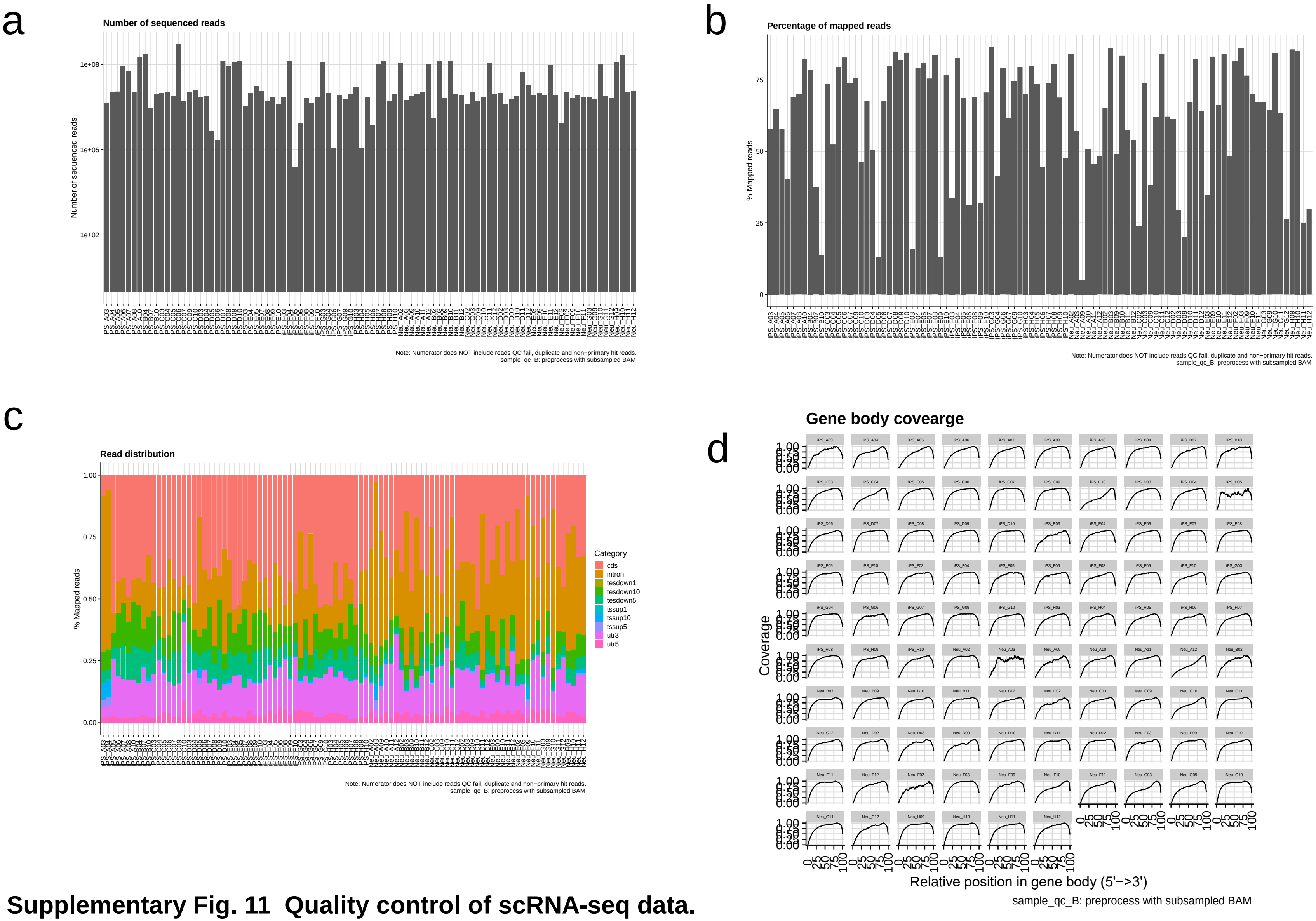


**Supplementary Fig. 11 Quality control of scRNA-seq data.**
(a) The number of total sequenced reads after the read trimming of FASTQ reads. (b) The percentage of reads mapped to the human genome. (c) Classification and distribution of uniquely mapped reads over genomic features. (d) Mean read coverage over transcripts.

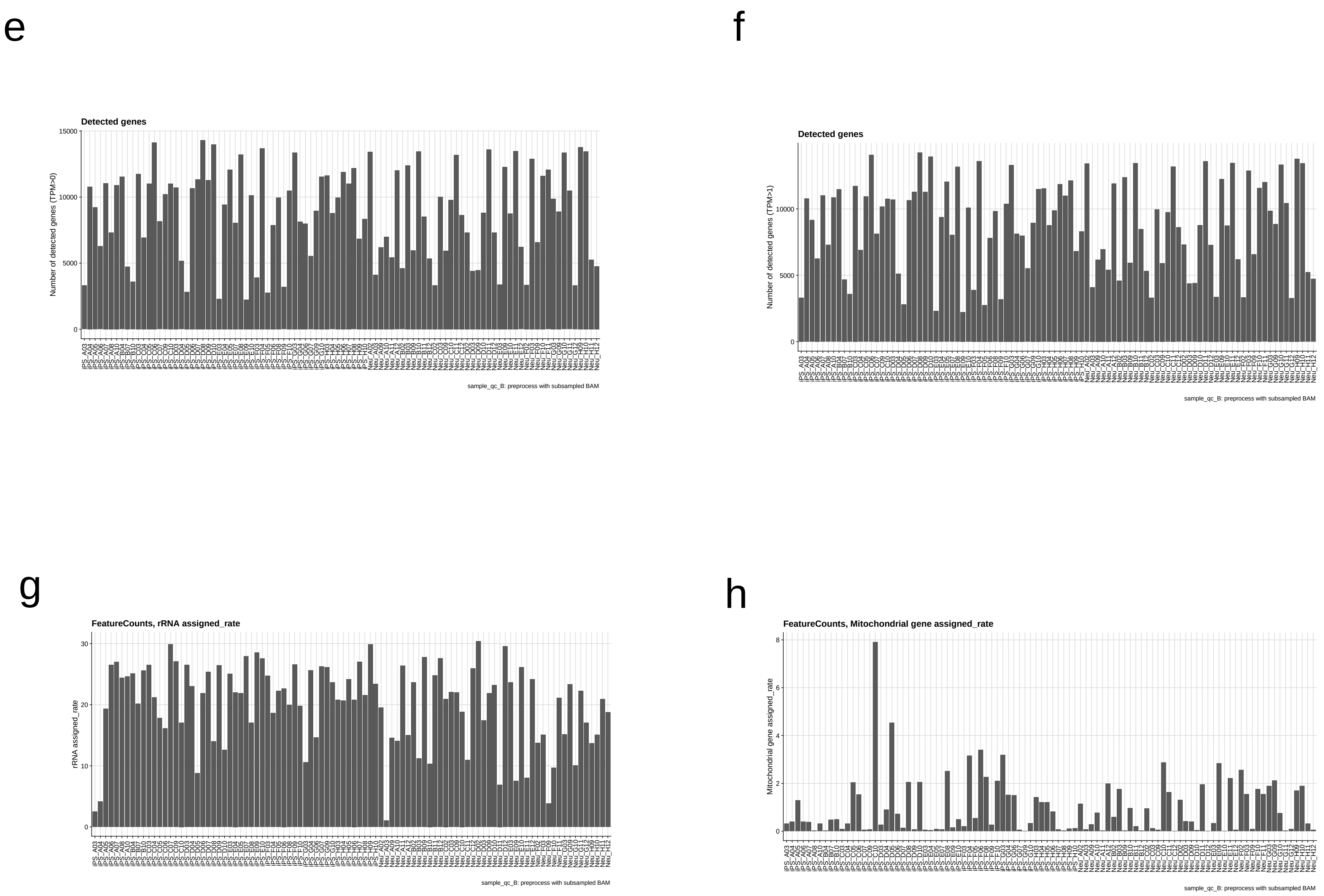


**Supplementary Fig. 11 Quality control of scRNA-seq data. (cont.)**
(e) The number of detected genes with TPM (transcript per million) > 0. (f) The number of detected genes with TPM (transcript per million) > 1. (g) The percentage of mapped reads that were overlapped with rRNA gene annotations on the genome. (h) The percentage of mapped reads that were overlapped with mitochondrial rRNA gene annotations on the genome.

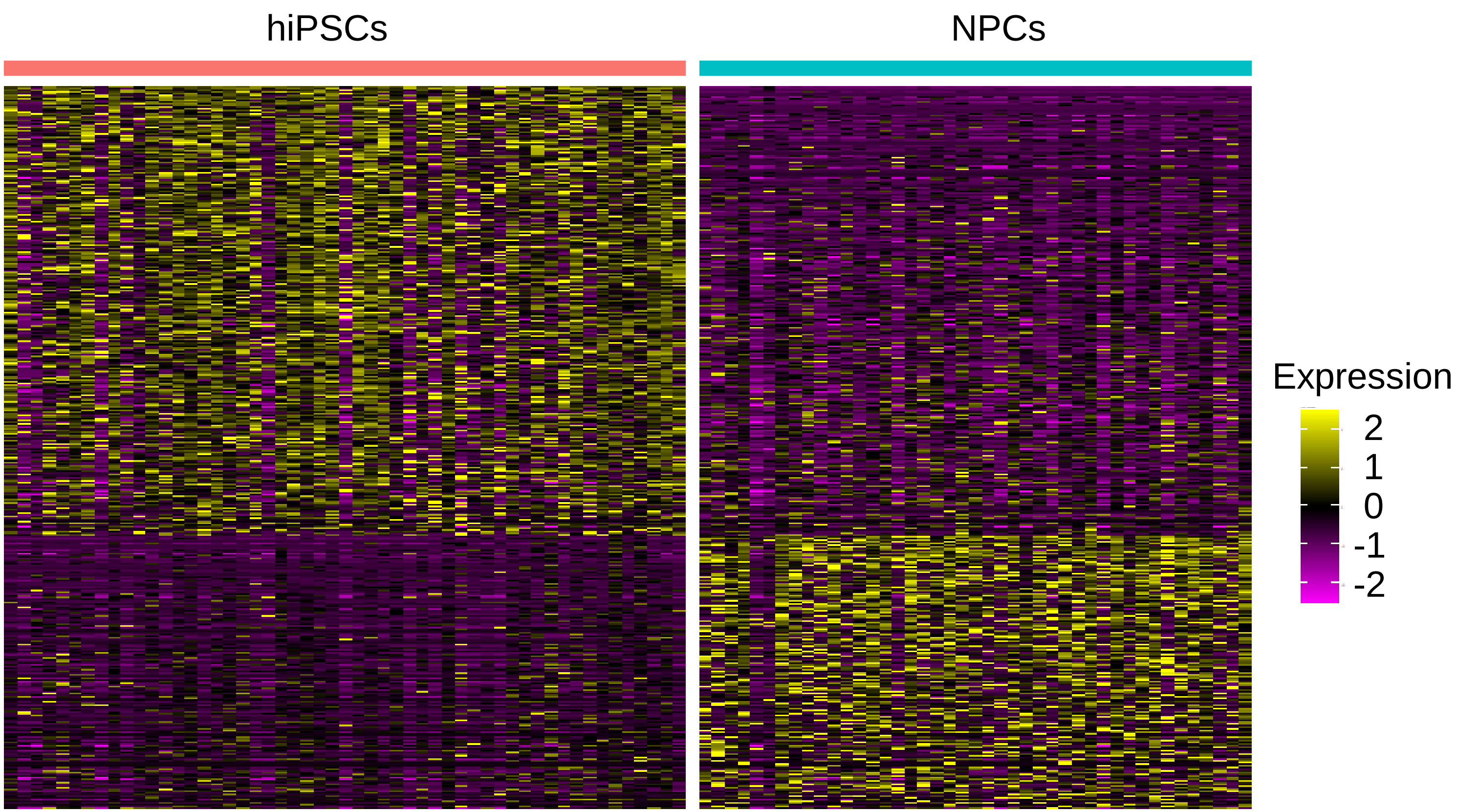


**Supplementary Fig. 12  scRNA-seq analysis of differentially expressed genes between hiPSCs and NPCs.** Heatmap of differentially expressed genes (DEGs) between iPSCs and NPCs. Criteria for DEG selection was set at log2(FoldChange) > 0.25 and Benjamini-Hochberg adjusted $p < 0.05$ (Mann-Whitney U test). Whole DEG lists were included in Supplementary Table 10.

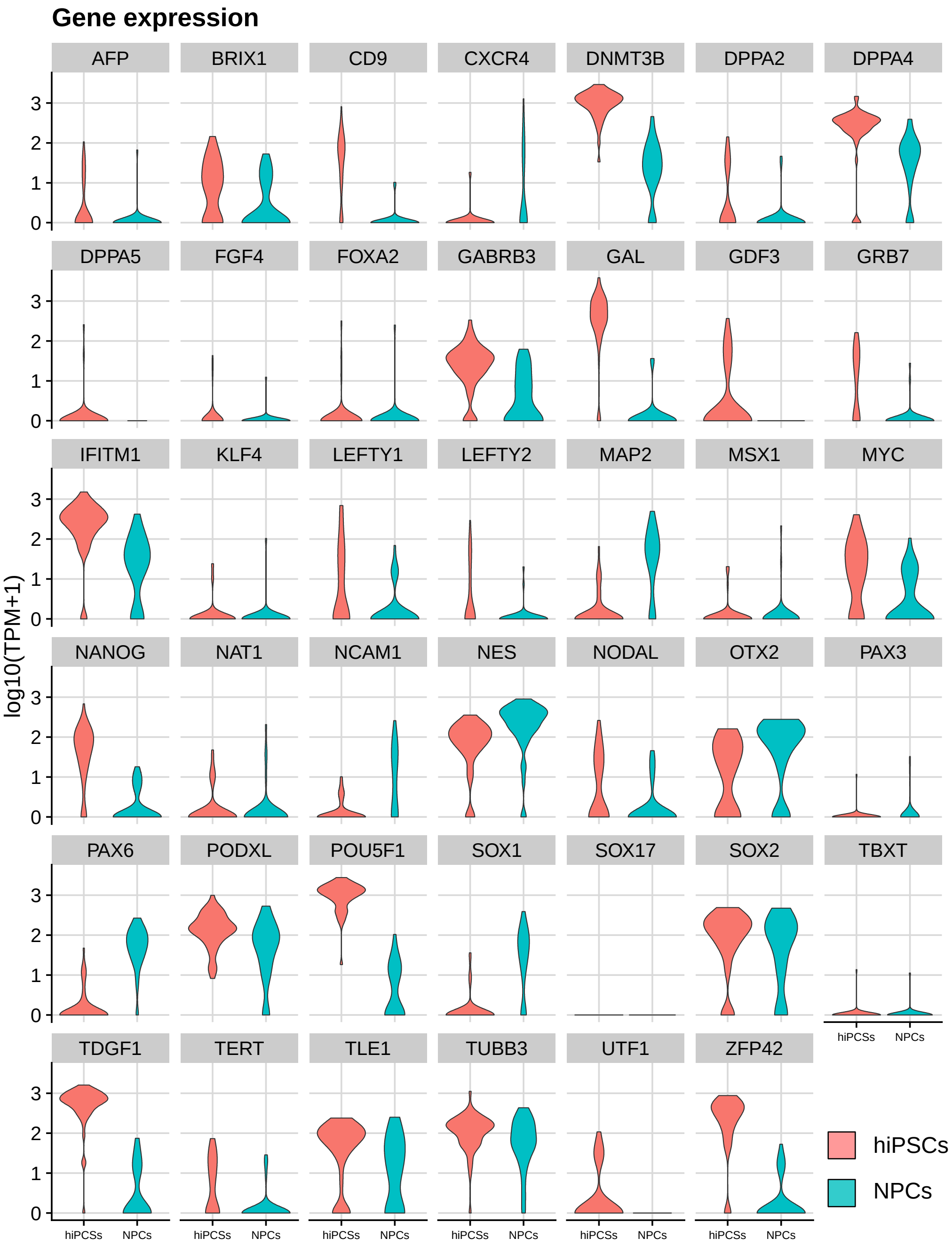


**Supplementary Fig.13 Gene expression of 41 selected markers by scRNA-seq.**
Violin plots show gene expression levels of 41 selected markers for hiPSCs (n = 53, *red*) and NPCs (n = 43, *cyan*) measured by scRNA-seq. Each point represents a single cell. The y-axes represent the log10(TPM+1).

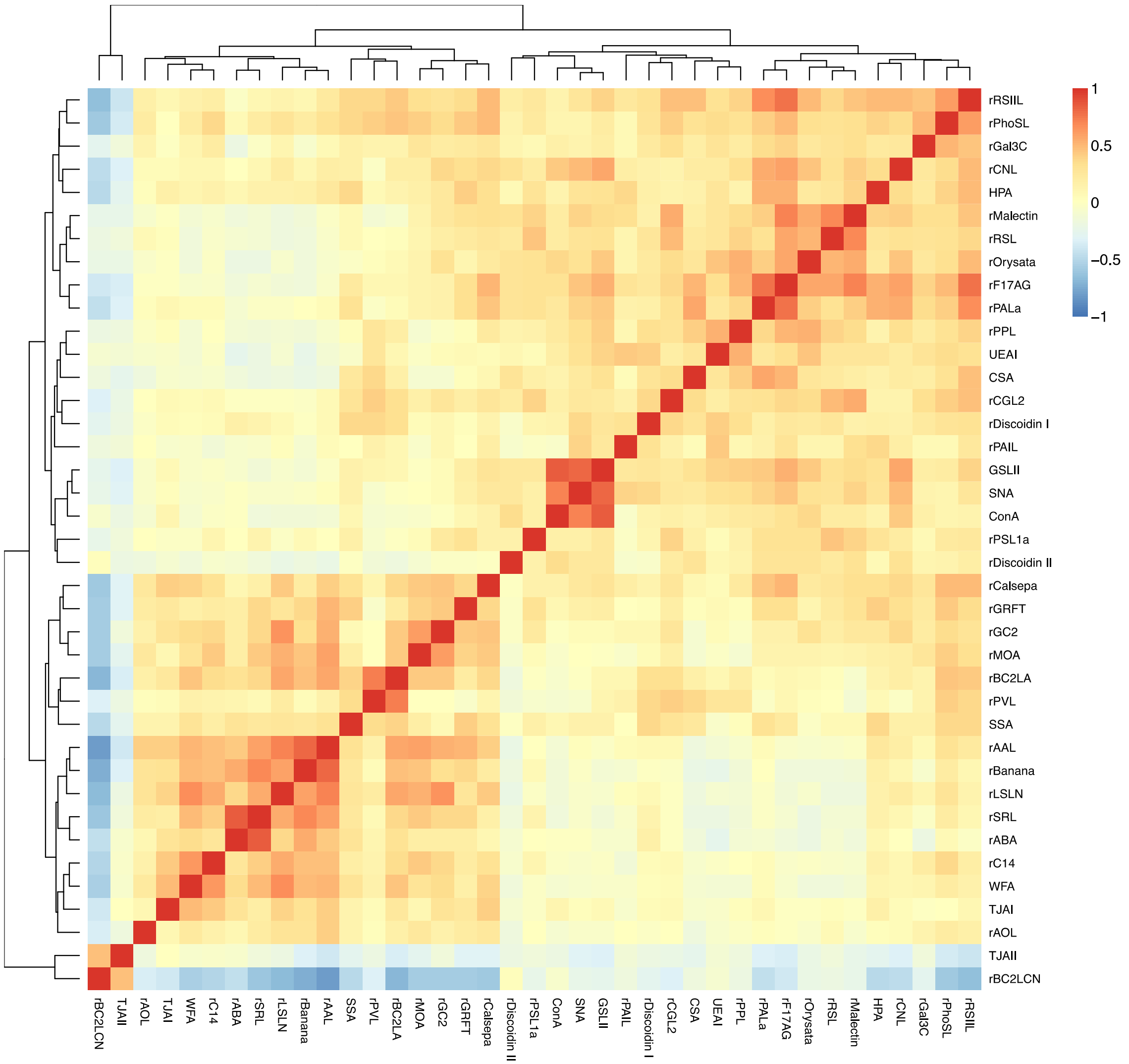


**Supplementary Fig. 14 Correlation of lectins across cells.**
A heat map shows the Pearson correlation coefficient of each pair of lectins. scGlycan-seq data of hiPSCs (n = 53) and NPCs (n = 43) were used. Rows and columns represent lectins.

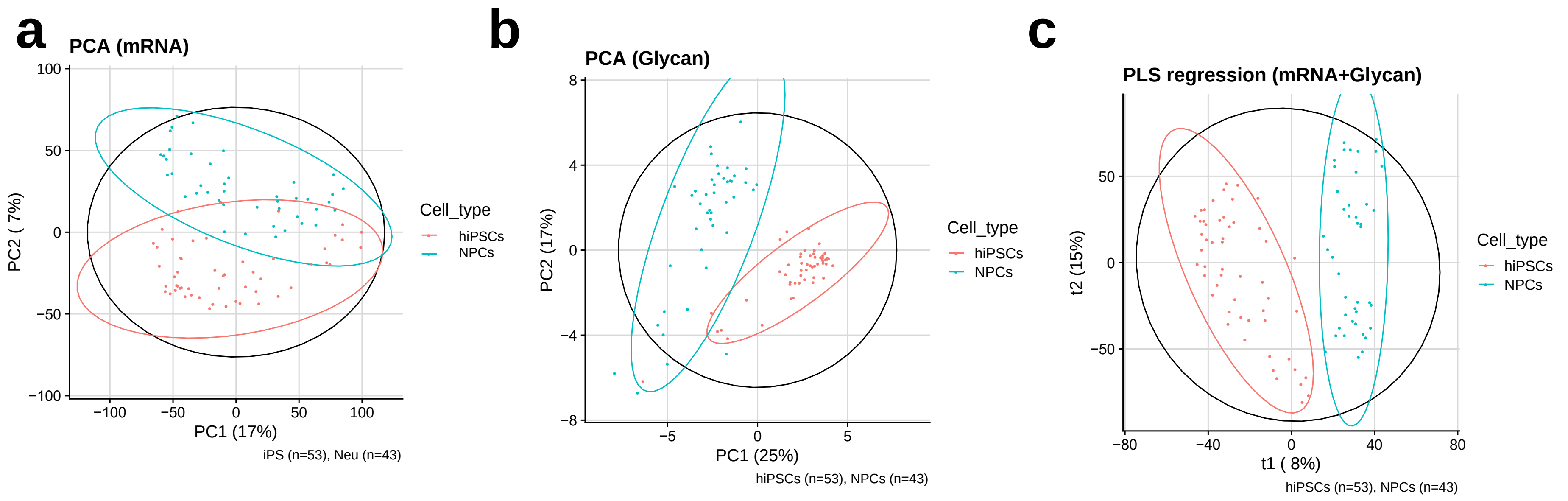


**Supplementary Fig. 15  PCA and PLS regression of scGR-seq.**
(a) PCA of scRNA-seq data of hiPSCs (n = 53, blue) and hiPSC-derived NPCs (n = 43, *red*). The x-axis and y-axis represent the first and second principal components (PC1 and PC2), respectively, and the percentages in the parenthesis indicate the proportions of explained variance. (b) PCA of scGlycan-seq data of hiPSCs (n = 53, *blue*) and hiPSC-derived NPCs (n = 43, red). The x-axis and y-axis represent the first and second principal components (PC1 and PC2), respectively, and the percentages in the parenthesis indicate the proportions of explained variance. (c) Partial least squares (PLS) regression analysis of scRNA-seq and scGlycan-seq data of hiPSCs (n = 53, blue) and hiPSC-derived NPCs (n = 43, red). The x-axis and y-axis represent the first and second components, respectively. Unlike PCA, PLS regression utilizes information from both scRNA-seq and scGlycan-seq data in the construction of components. scGlycan-seq data are available in Supplementary Tables 6, 7, 8, and 9.

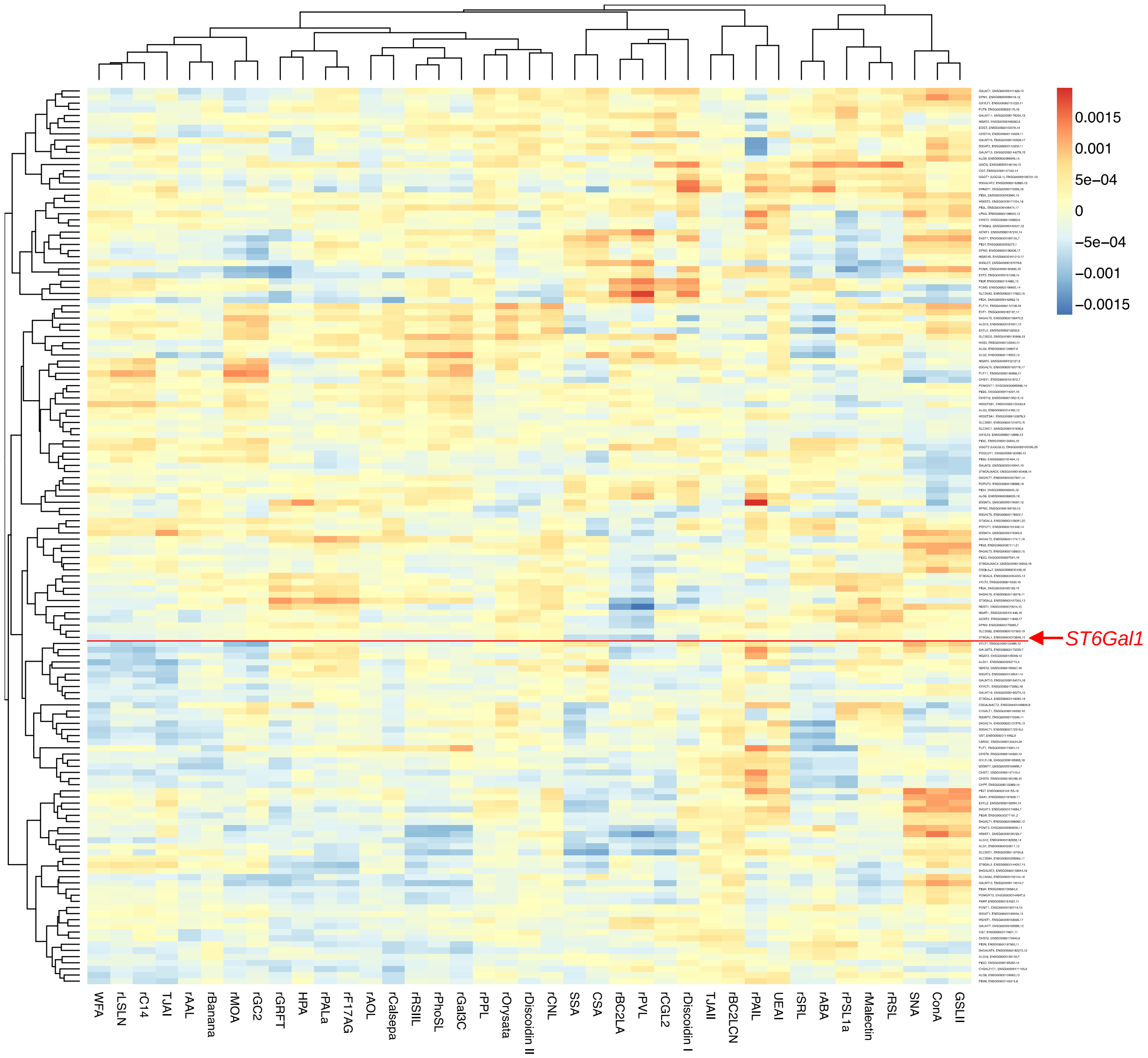


**Supplementary Fig.16 Correlation between lectin signal and glycosyltransferase gene expression.**
A heat map shows the association of glycogenes and lectins inferred by PLS regression. For each pair of a gene and a lection, the association was calculated by multiplying the gene loading and lectin loading for each component and summing the products across components. Rows represent genes and columns represent lectins.

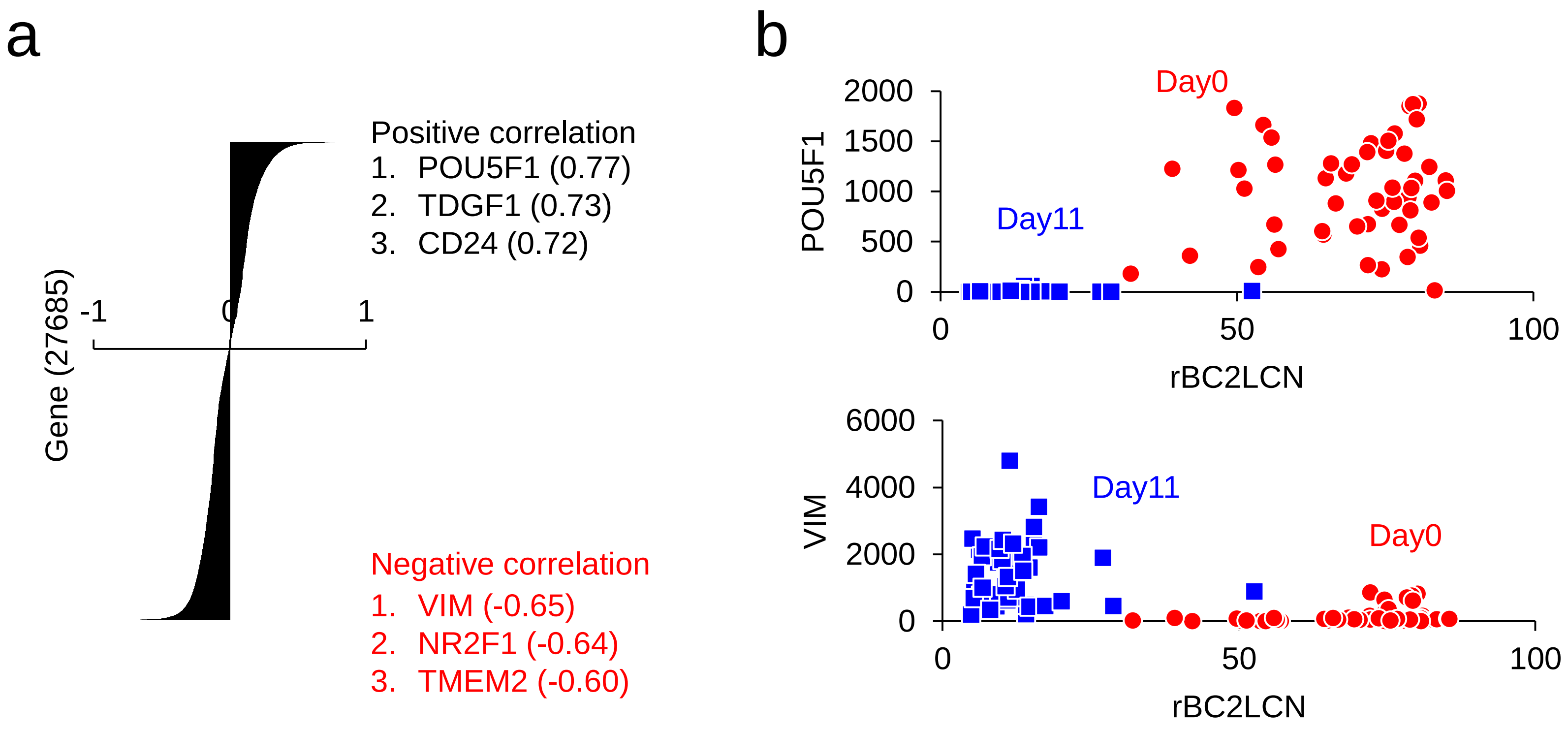


**Supplementary Fig.17 Correlation of genes with rBC2LCN in hiPSC and NPCs.**
(a) rBC2LCN showed the highest positive correlation with hPSC marker POU5F1 and the highest positive correlation with NPC marker VIM. (b) Correlation between rBC2LCN with *POU5F1* (*top panel*) and *VIM* (*bottom panel*).

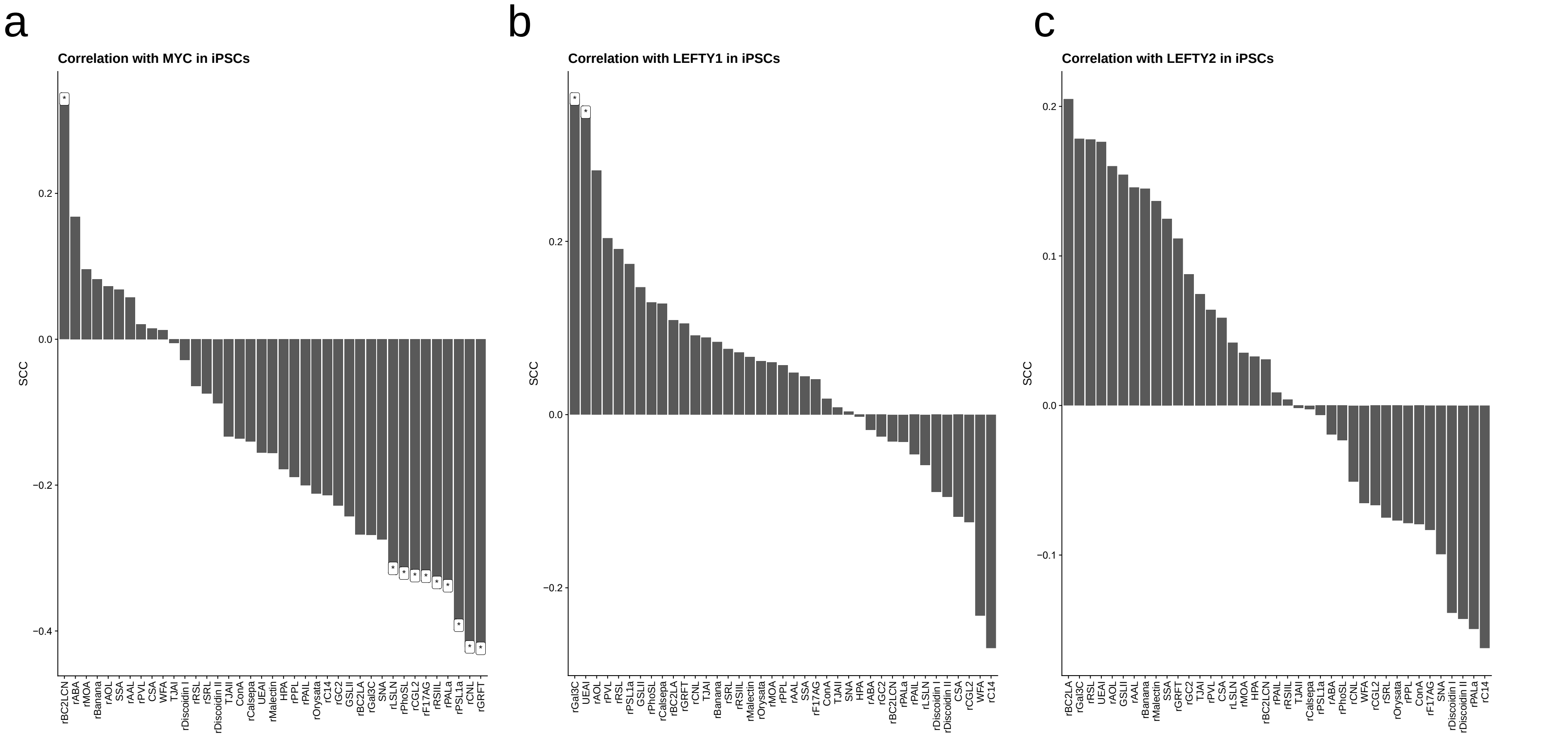


**Supplementary Fig.18 Correlation of lectins with fluctuating genes in hiPSCs.**
The Spearman's correlation coefficients were calculated between each lectin (scGlycan-seq) and each of MYC (a), LEFTY1 (b), and LEFTY2 (c) genes (scRNA-seq) across hiPSCs (n = 42). Cells with a low number of detected genes were removed. * $p < 0.05$
